# Tat-fimbriae (“tafi”) – novel type of haloarchaeal surface structures

**DOI:** 10.1101/2023.03.06.531322

**Authors:** Anna V. Galeva, Alexey S. Syutkin, Dahe Zhao, Igor I. Kireev, Alexey K. Surin, Elena Yu. Pavlova, Jingfang Liu, Hua Xiang, Mikhail G. Pyatibratov

## Abstract

In present study, we describe “**ta**t-**fi**mbriae (tafi)” – a novel type of archaeal surface appendages isolated from haloarchaeon *Haloarcula hispanica*. These **fi**lamental structures are unique because they are formed of protein subunits secreted through the **t**win-**a**rginine translocation pathway (Tat-pathway), in contrast to well-known archaeal surface filamentous structures secreted by the general secretory pathway (Sec-pathway). No cases of the role of Tat-pathway in the assembly of archaeal and bacterial filamentous structures have been described to date. “Tafi” are the first example of such structures. The precursor of the major tafi protein subunit TafA contains the N-terminal signal peptide carrying a twin-arginine consensus motif and fimbria-forming mature TafA lacks this signal peptide. We analyzed the gene neighborhood of the *tafA* homologues in the known haloarchaeal genomes and found a conservative cluster of seven associated genes *tafA, B, C, D, E, F, G*. We assume that all of them take part in the tafi synthesis. TafC and TafE proteins, whose precursor sequences also contain twin-arginine motifs, were detected as minor components of tafi. TafE protein is structurally similar to TafA, while TafC contains a TafA-like N-terminal domain and a C-terminal “laminin G-like” domain capable of functioning as an adhesin. TafD is annotated as a signal peptidase I. The functions of TafB, TafF and TafG are not known yet. This study demonstrated that Δ*tafA* and Δ*tafD* deletion mutant strains synthesized archaella and not tafi, and only tafi were detected in Δ*arlK* (gene of common archaellin/pilin signal peptidase) deletion strain. It was shown that the expression of complete *Har. hispanica taf*-gene cluster in a heterologous host *Haloferax volcanii* that does not have similar genes leads to synthesis of recombinant tafi structures similar to the native ones. The tafi function remains elusive, but our preliminary data suggest that these structures may be involved in cell adhesion to different surfaces or substrates.

## INTRODUCTION

The interest to study the third domain of life – Archaea – has increased in recent times. New species of Archaea are discovered with regularity. Many of them are extremophiles and exist in the most extreme environments on Earth, such as salt lakes, hot springs and places with extreme pH values. By now, Archaea have also been found in the ocean, soil, and even in the human gut and skin (Chaban *et al*., 2006; Mihajlovski *et al*., 2008; Baker *et al*., 2020).

Archaea have evolved different mechanisms for survival and adaptation to harsh environments. The surface structures of Archaea, such as archaella (previously *archaeal flagella*), pili, Iho670 fibers, Mth60 fimbriae, hami, cannulae, bindosomes etc. also perform an adaptive function. (Jarrell and Albers, 2012; Jarrell *et al*., 2013; Lassak *et al.*, 2012; Thoma *et al*., 2008). Among those structures, the most studied are pili and archaella, which structurally resemble bacterial type IV pili (T4P).

Archaella, similarly to bacterial flagella, enable cell motility. Using these organelles, cells can swim and move in their habitat. However, archaella and bacterial flagella fundamentally differ in their architecture, assembly mechanism, and origin: archaella share many common properties with bacterial T4P (Jarrell and Albers, 2012). In addition to their main role, archaella can mediate various functions such as adhesion to abiotic surfaces, formation of intercellular contacts, and, possibly, intercellular communication (Jarrell *et al*., 2021).

Archaeal pili discovered to date are structurally similar to type IV bacterial pili. They have been shown to be involved in adhesion, cell aggregation, biofilm formation and DNA transfer (Pohlschroder and Esquivel, 2015).

Archaeal adhesive pili (Aap) and UV-inducible pili (Ups pili) are the most studied pili in archaea. Ups pili are synthesized when the cells are exposed to UV rays. They were shown to be responsible for cellular aggregation, meaning that this process can help cells to exchange and repair DNA (Lassak *et al*., 2012).

Archaella and pili are widespread in Archaea, while other types of cell appendages are much less common. Specific surface structures (Iho670 fibers, Mth60 fimbriae, hami, cannulae, bindosomes, etc.) were found only in a few archaeal species and perform different functions such as adhesion to biotic and abiotic surfaces, cell-cell interaction, biofilm formation, etc. (Lassak *et al*., 2012). Their names are derived from the names of archaeal species or their structural and functional properties.

Iho670 fibers found in the archaeon of *Ignicoccus hospitalis* are involved in adhesion. They are composed of major protein Igni_0670 (Muller *et al*., 2009).

Hami were found in uncultivated euryarchaeal coccus SM1 living in cold (∼10°С) sulphidic springs as a part of archaeal-bacterial community. These filamentous structures carry a tripartite, barbed grappling hook at their distal end (the term “hamus” was proposed to describe it). Hami provide strong cell adhesion to various abiotic surfaces as well as to bacteria in their community (Moissl *et al*., 2005).

The lineament of *Pyrodictium* species is comprised of cells that grow in the network of extracellular tubules known as cannulae. These structures are highly resistant to high temperatures and denaturing agents such as 2% sodium dodecyl sulfate. The cryo–EM assay revealed that cannulae enters the periplasmic space, but not the cytoplasm of cells (Nickell *et al*., 2003).

Bindosomes were found in *Sulfolobus solfataricus*. It is assumed that these structures are integrated into the cell wall and facilitated the binding of saccharides from extracellular environment (Zolghadr *et al*., 2011).

The Mth60 fimbriae from *Methanothermobacter thermoautotrophicus* are composed of the 15 kDa Mth60 protein. Most likely, Mth60 is the only component of the fimbria, since it is able to form fibers similar to the native ones in vitro. Mth60 fimbria are about 5 nm in diameter and 10 µm in length. The significance of these structures for adhesion is proven by the fact that planktonic cells, as a rule, have only 1-2 fibres, while cells attached to the surface are covered with many fimbriae (Thoma *et al*., 2008).

Our research group have been studying the motility apparatus of an extremely halophilic archaea. These organisms thrive in environments with salt concentrations approaching saturation. One of the subjects of our research was the haloarchaeon *Haloarcula hispanica*. This is a model organism for studying haloarchaea, as it is easy to cultivate and genetically tractable. For genetic manipulation, we used the auxotrophic strain *Har. hispanica* DF60 with a deletion of *pyrF* gene (encodes the orotidine 5’-phosphate decarboxylase) unable to synthesize uracil, and plasmid pHAR for getting deletion mutants (Liu *et al*., 2011a

The present work describes filamentous structures, distinct from archaella and pili that we found in *Har. hispanica*. The main structural protein of these filaments TafA (HAH_0240) contains a motif used for twin-arginine translocation pathway (Tat-pathway) (Rose *et al*., 2002) instead of the signal peptide characteristic for the Sec-pathway that is typically used when exporting subunits of known archaeal filamentous structures. Therefore, we named detected structures tat-filaments (tafi). Tat-pathway is used for export and secretion of proteins in plants, bacteria and archaea. However, until now, the role of Tat-pathway in the assembly of filamentary structures has not yet been determined.

## RESULTS AND DISCUSSION

### *Haloarcula. hispanica* DF60 synthesize thin filamentous surface structures

Through the electron microscopy, we found that under certain conditions, *Har. hispanica* DF60 strain (with the deletion of *pyrF* gene involved in the pyrimidin synthesis) produced, along with typical archaella (10-11 nm in diameter), thinner filaments (about 3-5 nm in diameter) (**Figure 1**). Synthesis of these thin filaments also occurs in the wild type, but is much less noticeable than in DF60. In *Har. hispanica* DF60 cells thin filaments were observed extending from the surface of the cells. Both in cell samples and in preparations obtained by PEG precipitation (Gerl *et al*., 1989) thin filaments often formed bundles (**Figure 1**).

**Figure 1.**
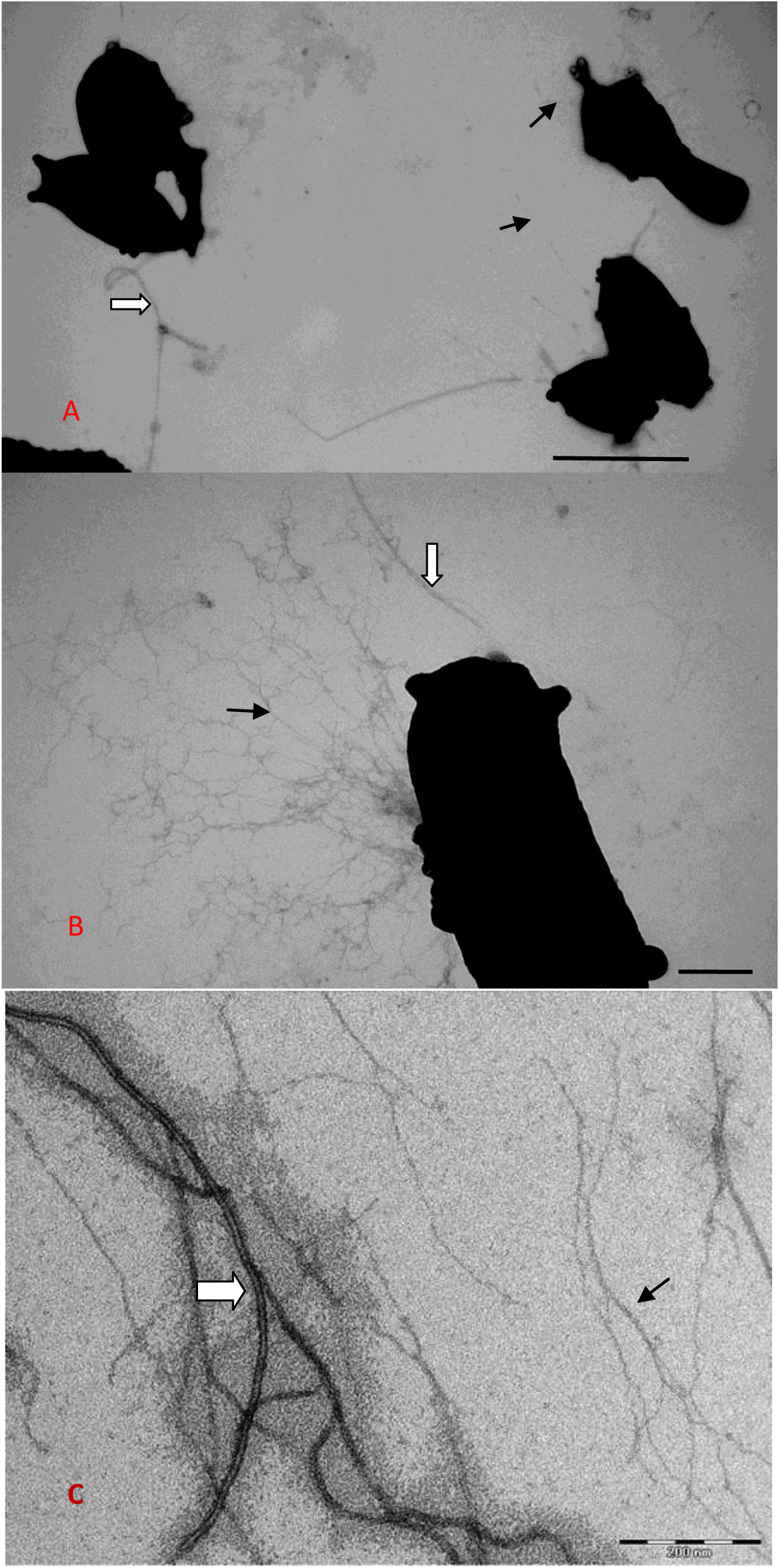
Transmission electron microscope micrographs of *Har. hispanica* DF60 cells (A, B) and surface structures isolated from *Har. hispanica* DF60 by PEG precipitation (C). White arrow points to the archaellum, black arrow points to the thin filaments. Scale bars are 200 nm (A, C) and 1 μm (B).

On the SDS-electropherogram, *Har. hispanica* archaellins look as a set of ladder-like bands. The intensity of these bands decays with decreasing mobility. Mass-spectrometric analysis of archaella samples prepared by high-speed centrifugation without PEG detected three archaellins ArlA1, ArlB and ArlA2 in each of the subbands of the “ladder”. This multiplicity is likely due to the presence of different archaellin glycoforms. In the samples taken after precipitation with PEG, in addition to the archaellin “ladder”, there was another major protein band observed on SDS-electropherogram; its relative proportion rose with an increase in the content of thin filaments in the preparations (**Figure 2**).

**Figure 2.**
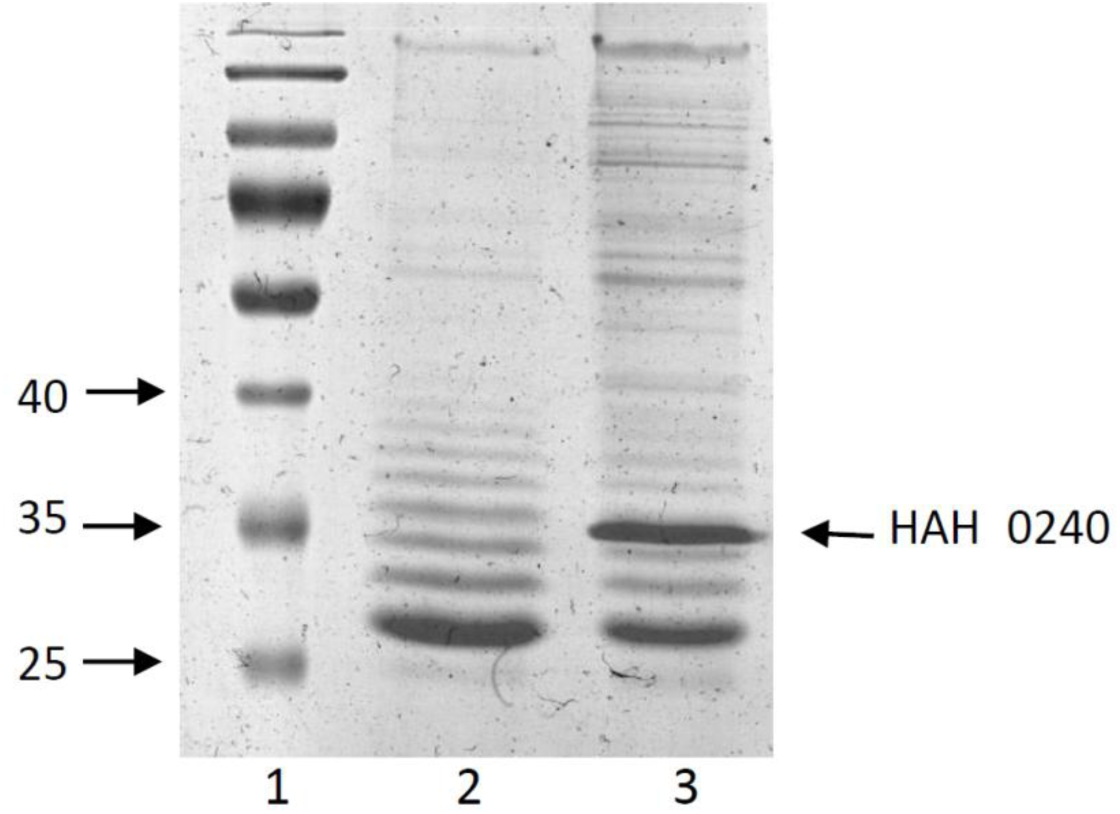
Samples of surface structures isolated from *Har. hispanica* strains analyzed at 14% SDS-PAGE. Lines: (1) Protein ladder (kDa); (2) Samples isolated from DF60 strain (Δ*pyrF*) using high-speed centrifugation without PEG and (3) using PEG precipitation (without prior high-speed centrifugation to separate archaella from thin filaments).

Mass-spectrometry revealed that the extra protein band in DF60 strain sample corresponds to the WP_044951594.1 (HAH_0240 or HAH_RS01150) annotated as a hypothetical *Har. hispanica* protein (we will refer to it as TafA) (**Figure 3, Figure 1S**).

**Figure 3.**
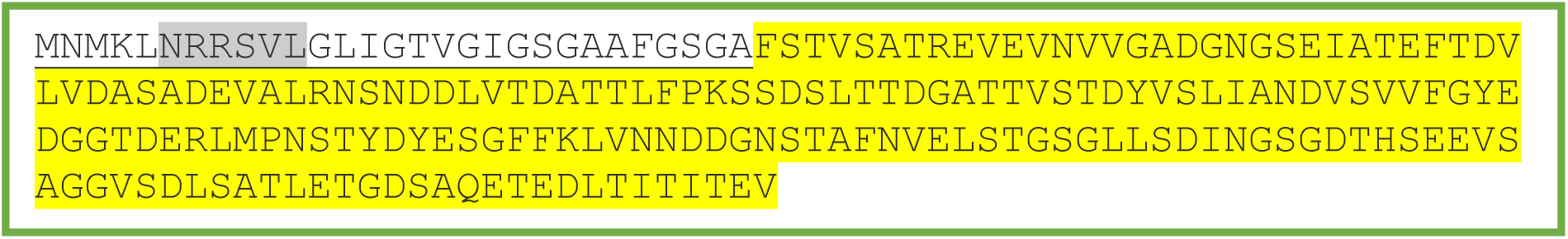
Amino acid sequence of *Har. hispanica* TafA (HAH_0240) precursor protein. The Tat pattern is highlighted in grey. The mature protein sequence is highlighted in yellow.

### Structural features of TafA (HAH_0240) protein

Gene *tafA* encodes a sequence of 208 or 210 amino acid residues (it depends on which of the two neighboring methionine codons will be the start). Its sequence does not show noticeable homology with known proteins, including those of other filamentous structures such as archaellins and pilins. A signal peptide necessary for secretion by the Sec-pathway is not detected in it. Surprisingly, using the Tat-find algorithm (Rose *et al*., 2002) we found the pattern for the twin-arginine translocation pathway (NRRSVL) (**Figure 3, Table 2S**).

The twin-arginine translocation pathway (Tat-pathway) is an alternative to the Sec-pathway. The main difference between these two protein secretion pathways is that the substrates for the Sec-pathway are transported in an unfolded form, whereas substrates for the Tat-pathway are folded before they are transferred across the membrane (Szabo and Pohlschroder, 2012). Tat-system obtained its name from the fact that their substrate sequences must contain two conserved adjacent arginines in the signal peptide. As mentioned above, all archaeal and bacterial surface filamentous structures known to date are formed using the Sec-pathway. Thus, we hereby describe the first type of prokaryotic filamentous surface appendages which depend on the Tat-system. These structures are therefore called “tat-fimbriae - tafi”.

The SignalP 5.0 server (Almagro Armenteros *et al*., 2019; https://services.healthtech.dtu.dk/service.php?SignalP-5.0) predicted the cleavage site of the signal sequence of TafA protein to be between 35 (alanine) and 36 (threonine) amino acids. Using mass spectrometry we found N-terminus of mature TafA to be between 29 (alanine) and 30 (phenylalanine) amino acids (**Figure 3, Figure 1S, Table 2S**).

Apparently, not only the secretion mechanism but also the structure of the TafA protein significantly differs from known archaeal pilins and archaellins (archaeal flagellins). As we know, the structure of type IV pili and archaella is supported by hydrophobic interactions between the N-terminal α-helices of archaellins and pilins. According to the theoretical prediction of the secondary structure by PsiPred program (Buchan *et al*., 2013), the mature TafA protein contains only unstructured regions and β-strands (**Figure 2S**).

In addition, we discovered that tafi could completely dissociate at NaCl concentrations below 5% and while heating the solution to 90°С. In these conditions, TafA protein is likely to become converted into a monomeric form (according to data of gel filtration chromatography). Circular dichroism measurements confirmed that TafA protein was in an unfolded state at 0% and 5% NaCl concentrations. With a further increasing of salt concentration, we observed β-structure spectra characteristic (**Figure 4**). These findings are consistent with the theoretical predictions for the TafA secondary structure.

**Figure 4.**
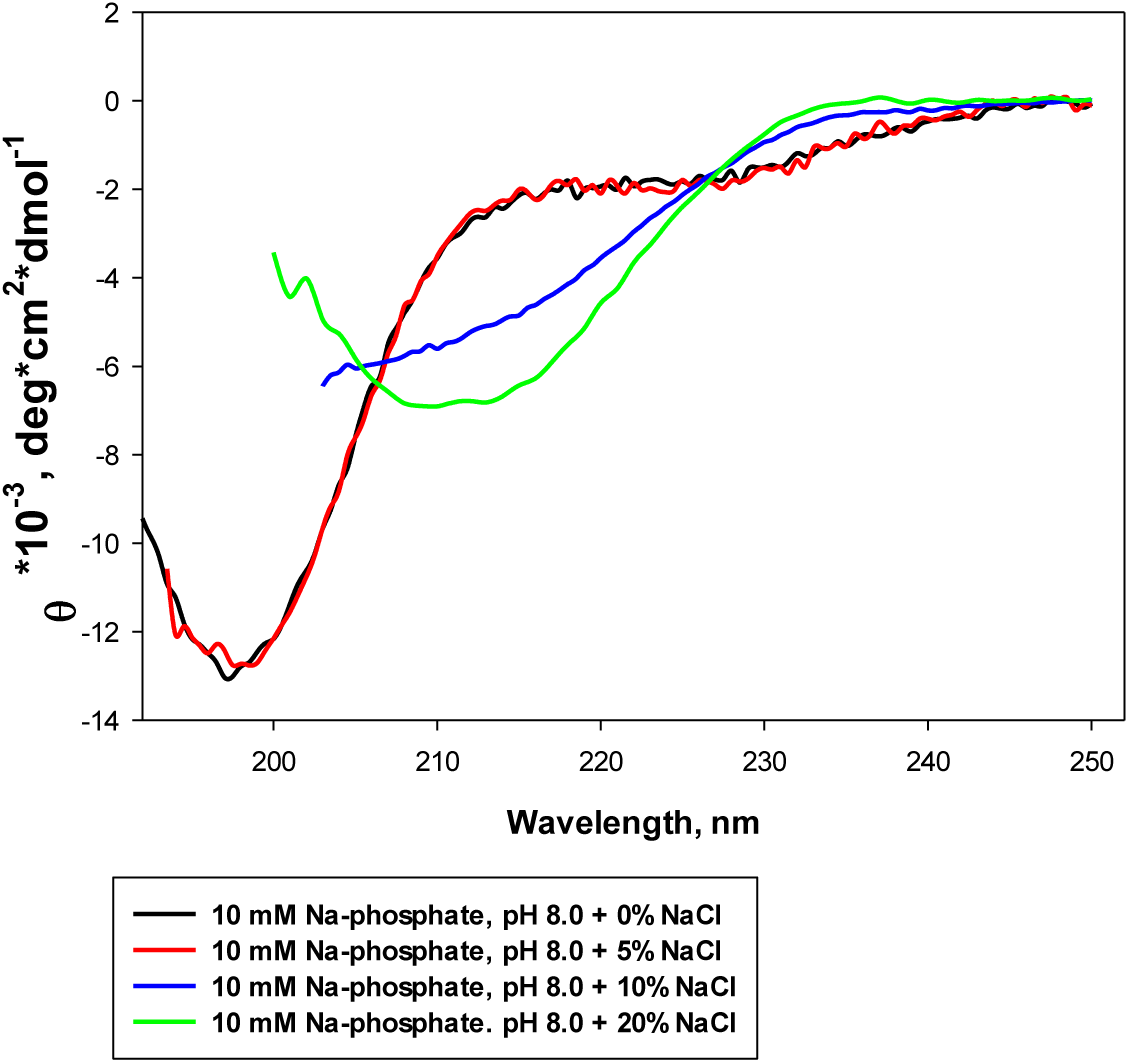
CD spectra of isolated TafA protein at different salt concentrations.

Microcalorimetric studies confirmed that at salt concentrations corresponding to native (20% - 25% NaCl) monomeric protein TafA has a domain structure (a single heat absorption peak was observed at a melting point of about 72 °C, when melting is reversible). However, dissociated TafA was unable to spontaneously polymerize into fimbria resembling natural ones. In vitro TafA polymerization experiments conducted under various conditions gave a negative result (data not shown).

### Deletion of *tafA* and *tafD* genes stops the tafi synthesis

As mentioned above, along with archaella we found a new type of filamentous structures in *Har. hispanica* DF60. To confirm our proposition that TafA protein is the main structural component of these filaments, we deleted the *tafA* (*hah_0240*) gene as described in the experimental procedure. We could not detect the protein TafA band on the SDS-PAGE (**Figure 5**) in the samples prepared from Δ*tafA* cells, and the corresponding filamentous structures were not found using electron microscopy (**Figure 6**). The loss of the ability of Δ*tafA* mutant cells to synthesize tafi confirms our suggestion about TafA protein being the main structural component of tat-fimbria.

**Figure 5.**
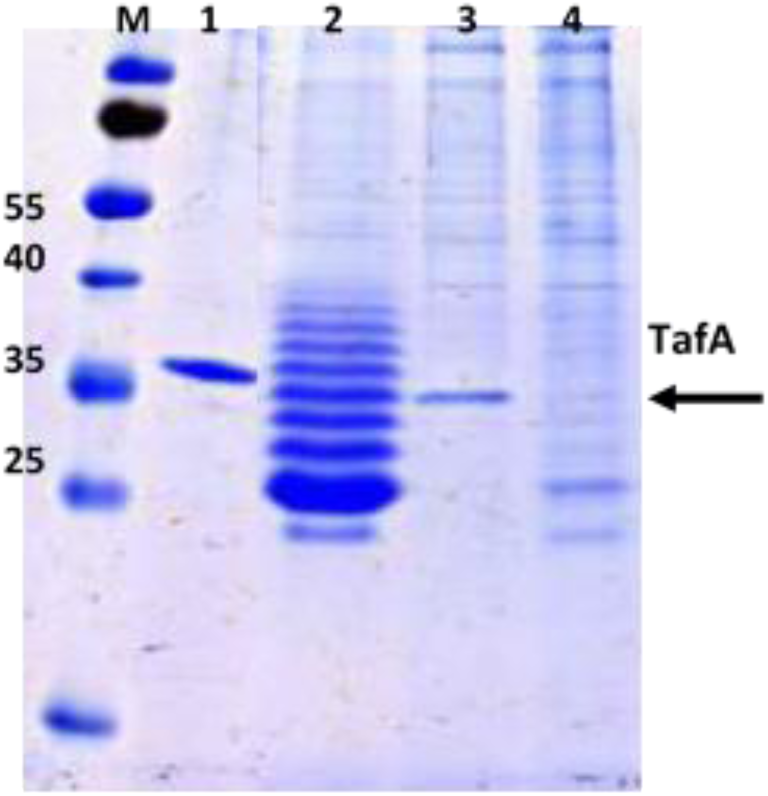
SDS-electropherogram of of purified TafA protein (1) and surface structures isolated from DF60 strain (1), Δ*hah_0243* (*tafD*) (2), Δ*arlK* (3), Δhah_0240 (*tafA*) (4). M – protein ladder.

**Figure 6.**
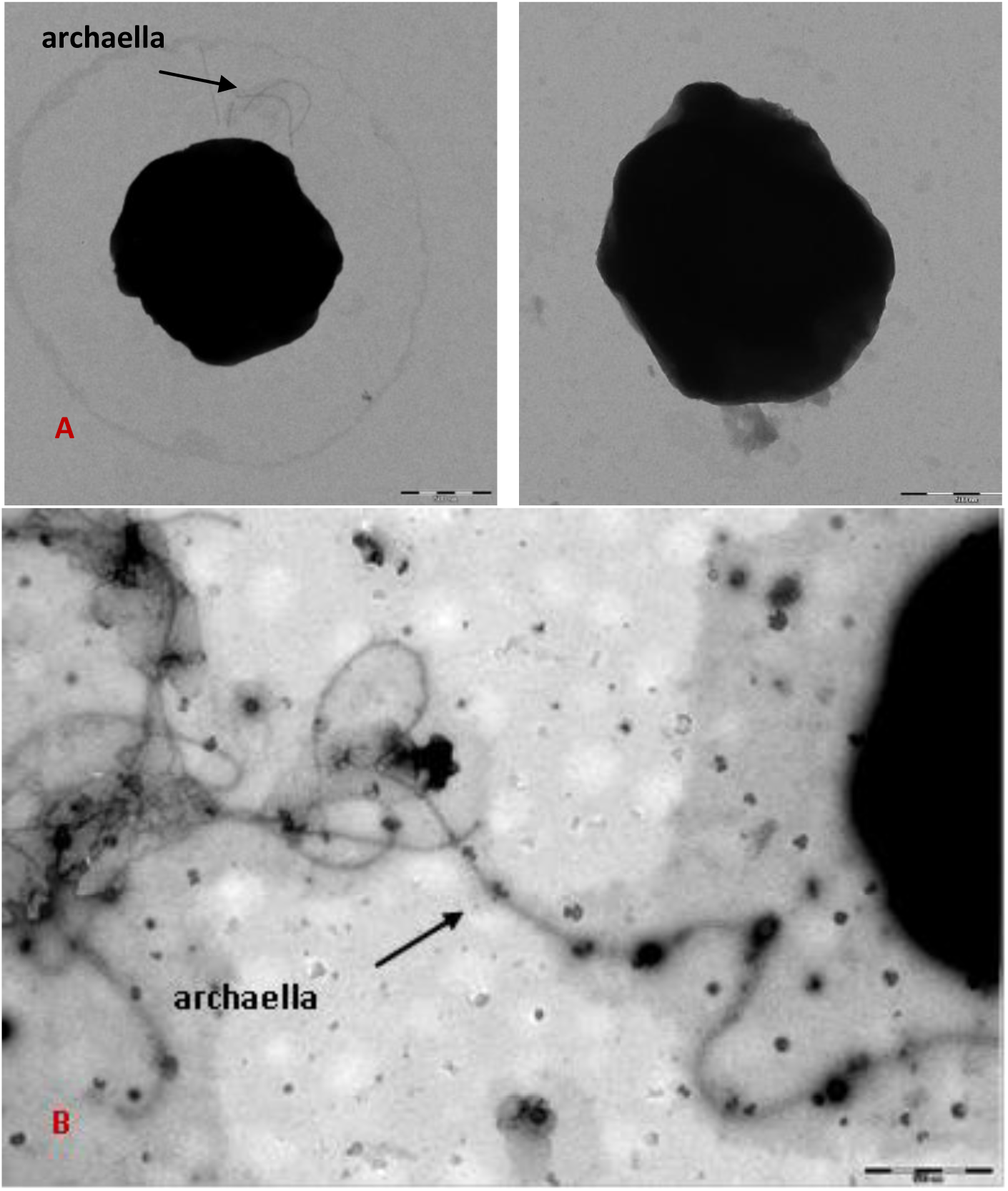
Electron microscopic micrographs of cells (A) and surface structures (B) isolated from Δ*tafA* strain. Scale bar is 500 nm.

### Deletion of *arlK* gene

For detailed study of tafi, it would be useful to obtain a strain without archaella and pili. To do this, we constructed a strain with deletion of *arlK* (*hah_2955*) gene encoding a common signal peptidase of archaellins and pilins (ArlK). As shown by our experimental data, the cells of Δ*arlK* strain contained only long thin fimbria and did not contain archaella and pili. In the samples from these cells, we detected the protein band of TafA, but the archaellin bands were lost (**Figure 5**). Electron microscopy data confirmed that tafi are localized on the cell surface and somehow are anchored into it (**Figure 7**).

**Figure 7.**
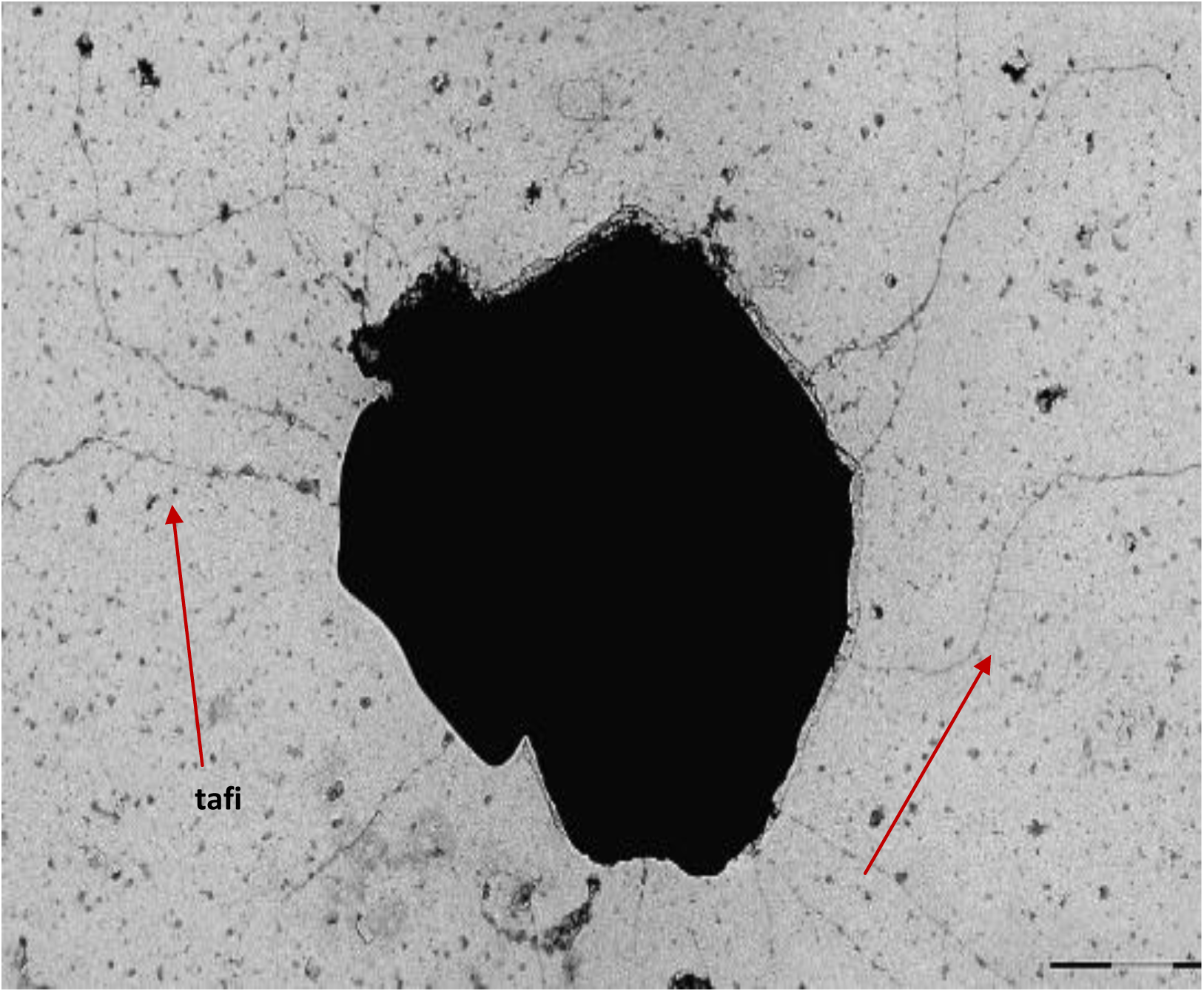
Electron microscopic micrograph of cells from Δ*arlK* strain. There are only tafi structures extending from the cell body. Scale bar is 1 μm.

### Heterologous expression of Har. hispanica tafA-tafG genes in Haloferax volcanii

Previously, it was shown that functional archaellar filaments are able to assemble upon the expression of *Halorubrum lacusprofundi* or *Halorubrum saccarovorum* archaellin genes in a heterologous system of a non-motile *Hfx. volcanii* strain and restore its motility (Pyatibratov *et al*., 2020). For similar experiments, it is convenient to use a special strain of *Hfx. volcanii* MT78 in which the archaellin operon (Δ*arlA1*Δ*arlA2*) and the genes required for pili assembly (Δ*pilB3*-*C3*) were deleted (Esquivel *et al*., 2014). Thus, this strain is non-motile and does not have any structures related to type IV pili. We decided to use this system to test the possibility of tafi synthesis in a heterologous system. Remarkably, the *Hfx. volcanii* genome, unlike its closest relatives *Hfx. elongans*, *Hfx. larsenii* and *Hfx. mediterranei*, does not contain its own genes corresponding to *Har. hispanica tafA-tafG* genes. Therefore, we suggested that genes of tafi proteins could be successfully expressed in the *Hfx. volcanii* system.

For this purpose, a vector pAS24 containing gene set from *tafA* to *tafD* was constructed based on pTA1228. In this vector, *tafA* gene was positioned under a tryptophan-induced promoter, and *tafB*, *tafC* and *tafD* genes were under their natural promoters (**Figure 3S**). This set includes the genes for the main tafi protein TafA and TafD, a putative signal peptidase that processes the TafA precursor. In the case of the native host *Har. hispanica*, we showed that both of these proteins were important for tafi synthesis. However, during the expression of the pAS24 plasmid in *Hfx. volcanii* we did not observe the formation of structures resembling tafi. We hypothesized that the translocation of TafA is impaired, and it either accumulates in the cell or is secreted but cannot form fimbria.

Experimental data confirmed that TafA is secreted but not polymerized. The supernatant obtained after removing the cell biomass after several rounds of centrifugation was concentrated by 100 times with Amicon Ultra Filtres. The collected product was analyzed by SDS-PAGE. As a control, we used the same preparation from MT78 Hfx. volcanii without expression vector. As seen in **Figure 8**, the concentrated supernatant of MT78/pAS24 contains four additional protein bands.

**Figure 8.**
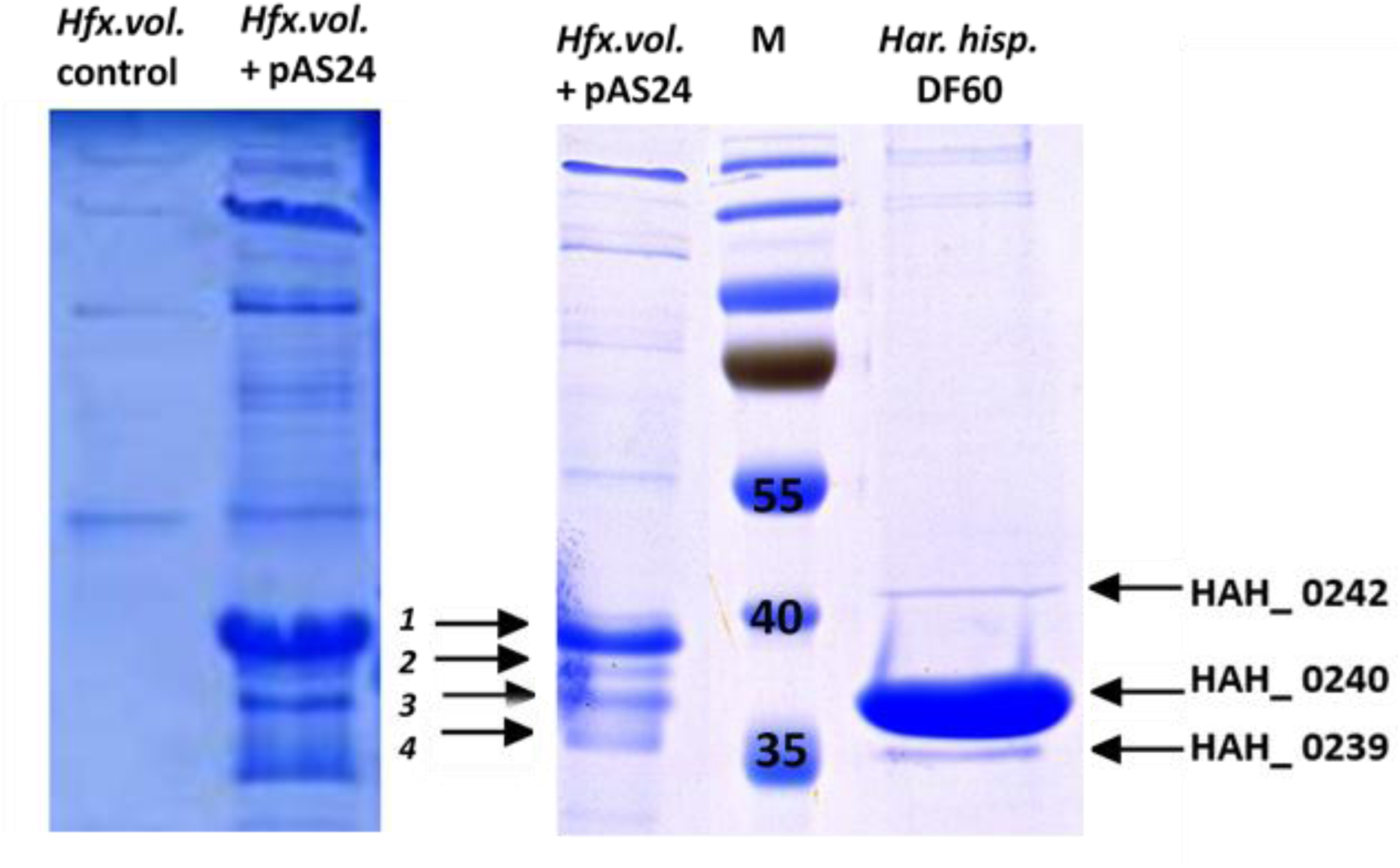
SDS-electropherogram of the heterologous expression of pAS24 plasmid (with incomplete set of *Har. hispanica taf* genes (*tafA* - *tafD*) in *Hfx. volcanii*.

Mass-spectrometry revealed that all 4 protein bands contained protein products of *tafA* (*hah_0240*) and *tafC* (*hah_0242*) genes (**Figure 4S**). TafA and TafC proteins were identified with the highest confidence (87% and 55% coverage) in bands 3 and 1, respectively. Both TafA and TafC contain the twin-arginine patterns. This indicates that both of these proteins (TafA and TafC) successfully used the Tat-pathway for export and TafD (presumable signal peptidase) processed them. Surprisingly, in all bands, along with TafA and TafC proteins, the *Hfx. volcanii* HVO_A0133 (WP_013035207.1) protein was detected (the largest coverage of 83% was obtained for band 4). It is annotated as glycine zipper family protein and contains a Tat-pathway signal sequence. This protein is encoded by a plasmid pHV4 and is not universal; its close homologues were found in only two haloarchaea (*Halorubrum* sp. Atlit-28R and *Halomicrobium katesii*). The function of HVO_A0133 and the effect of pAS24 plasmid expression on its synthesis remain unclear. The observed heterogeneity of proteins TafA, TafC and HVO_A0133 may be associated with probable variability in their processing. These results show that TafA and TafC can be synthesized and secreted in the *Hfx. volcanii* heterologous system; however, the recombinant TafA protein cannot polymerize and form tafi. We suspected that inability of TafA protein to polymerize in these circumstances may be caused by an absence of additional (adaptor) proteins encoded by other taf-genes (*tafE*, *F*, *G*). Therefore, our next step was to obtain an extended genetic construct pAS30, based on pAS24, containing the full set of genes *tafA* – *tafG* (**Figure 5S**). We showed that the expression of the full set of seven taf-genes in the plasmid pAS30 in the *Hfx. volcanii* resulted in the synthesis of recombinant tat-fimbria similar to natural ones from *Har. hispanica* (**Figure 9**). These results confirmed that the polymerization of the main tafi subunit TafA requires the presence of accessory proteins encoded by the taf-cluster. Future experiments will be needed to clarify the role of each of these genes in the assembly and function of these fimbriae.

**Figure 9.**
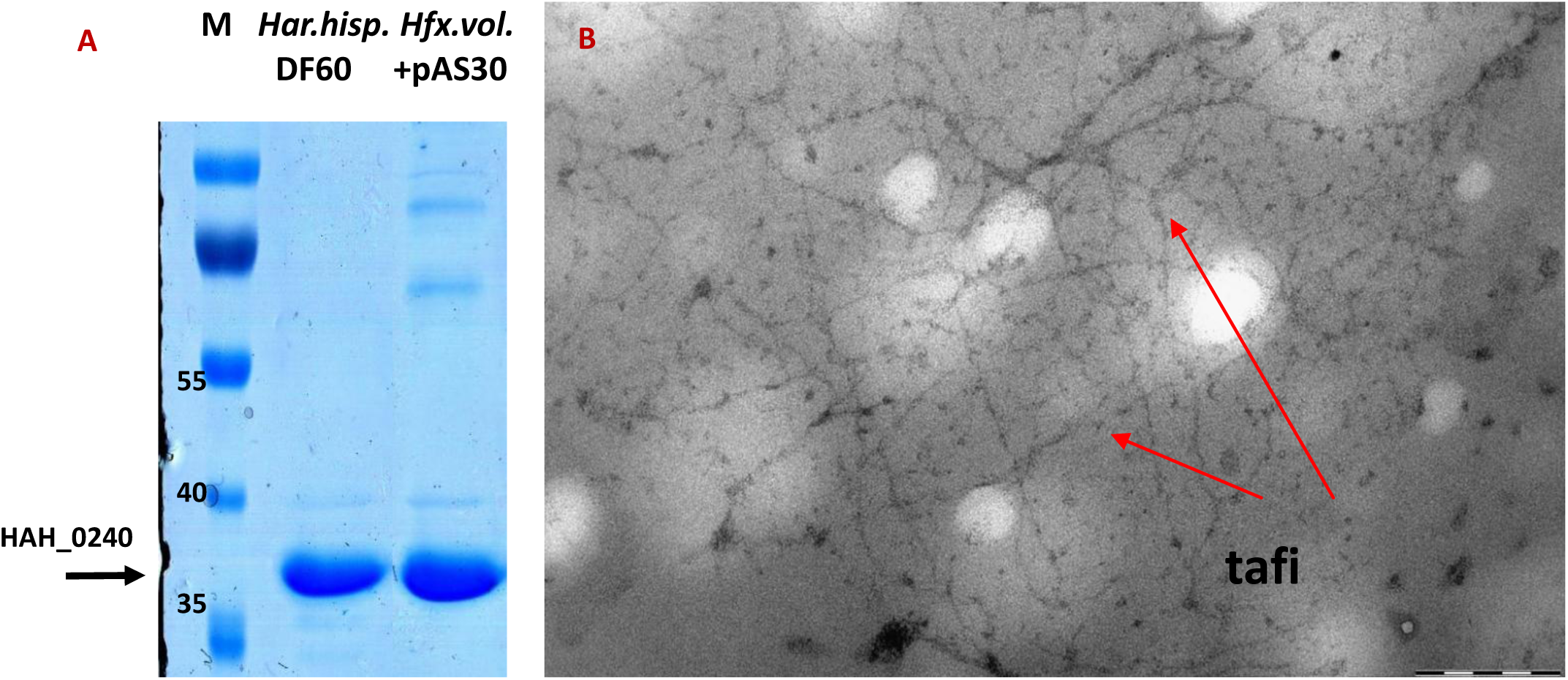
The heterologous expression of pAS30 plasmid (with the full set of *Har. hispanica taf* genes (*tafA* – *tafG*) in *Hfx. volcanii*: 16% SDS-PAGE (A) and electron microscope micrograph of recombinant tafi structures, scale bar is 200 nm (B).

To confirm the key role of the Tat-pathway in TafA secretion followed by assembly into tafi, we performed nucleotide substitution mutagenesis of pAS30 plasmid to recode the arginine pair – the key to the Tat pathway of TafA – to a lysine pair. The modified pAS31 plasmid was used to transform cells of *Hfx. volcanii* MT78. Using the technique that was effective for isolating tafi from the MT78/pAS30 strain, we obtained similar preparations from the MT78/pAS31 strain. However, we were unable to detect any filamentous structures in these preparations. Thus, we have confirmed that the replacement of RR by KK is critical for the assembly of tafi (data not shown).

### Bioinformatical analysis of *tafA* gene and surrounding

Currently, the product of *tafA* gene is annotated as “hypothetical protein”. This gene is surrounded by ORFs, which also encode “hypothetical proteins”, and does not seem to be in one operon with them. Using the BLAST algorithm (http://blast.ncbi.nlm.nih.gov/Blast.cgi), we found that TafA protein is not universal and is limited to halophilic archaea. Explicit sequence homologues were found in many representatives of the genera *Haloarcula*, *Haloferax*, *Halorubrum* and in several members of other genera: *Halorhabdus* sp. CBA1104, *Haloterrigena mahii* H13, *Natrialba* sp. INN-245 *Natrinema* sp. SLN56, *Natronorubrum thiooxidans* HArc-T and *Salinadaptatus halalkaliphilus* XQ-INN 246 (**Table 1S**).

At this rate, the closest homologues (100% to 67% identity) of TafA protein have been found in approximately 20 known genomes, including *Haloarcula amylolytica* JCM 13557, several *Har. hispanica* strains, *Har. rubripromontorii* pws8, *Har. rubripromontorii* SL3, *Har.* sp. Atlit-7R, *Har.* sp. Atlit-47R, *Har.* sp. Atlit-120R, *Har.* sp. CBA1115, *Har.* sp. CBA1127, *Har.* sp. CBA1128, *Har.* sp. CBA1131, *Har.* sp. K1, *Har. taiwanensis*, *Haloferax elongans* ATCC BAA-1513, *Hfx. larsenii* CDM_5, *Hfx. larsenii* JCM 13917 and *Hfx. mediterranei* ATCC 33500. The genomes of the second group of haloarchaea (*Haloarcula argentinensis, Har. californiae, Har. marismortui, Har. sinaiiensis, Har. vallismortis, Halorubrum coriense, Hrr. ezzemoulense, Hrr. hochstenium, Hrr. litoreum*, etc.) also contain genes encoding similar proteins with significantly less homology (about 40% identity) (**Table 1S**). In addition, the genomes of many halophilic archaea encode proteins in which only the first 40 N-terminal amino acid residues (including the signal peptide) are homologous to the TafA protein, and the central and C-terminal parts do not have any homology.

A more sophisticated assay of *Har. hispanica* genome shows that it contains a cluster of seven genes that we believe to be involved in tafi synthesis. We named them taf genes (identification marks are given at **Figure 10** and **Figure 6S**). Most of the encoded proteins in this cluster (except signal peptidase I, TafD) are annotated as hypothetical. According with preliminary survey (with TatFind and SignalP-5.0 servers), the HAH_0239 (TafE) and HAH_0242 (TafC) proteins contain twin-arginine patterns and appear to be translocated via Tat-pathway (**Table 2S**). On SDS-electropherograms of comparatively pure tafi preparations, two minor bands can be seen above and below the major band of the TafA. Mass-spectrometric analysis showed that the upper band corresponds to the TafC, and the lower band to the TafE (**Figure 8, Figure 7S**). Based on the results of scanning the gels, we estimated the approximate molar ratio between these proteins in filament preparations: for every 100 molecules of TafA, there is approximately one TafC molecule and five TafE molecules.

**Figure 10.**
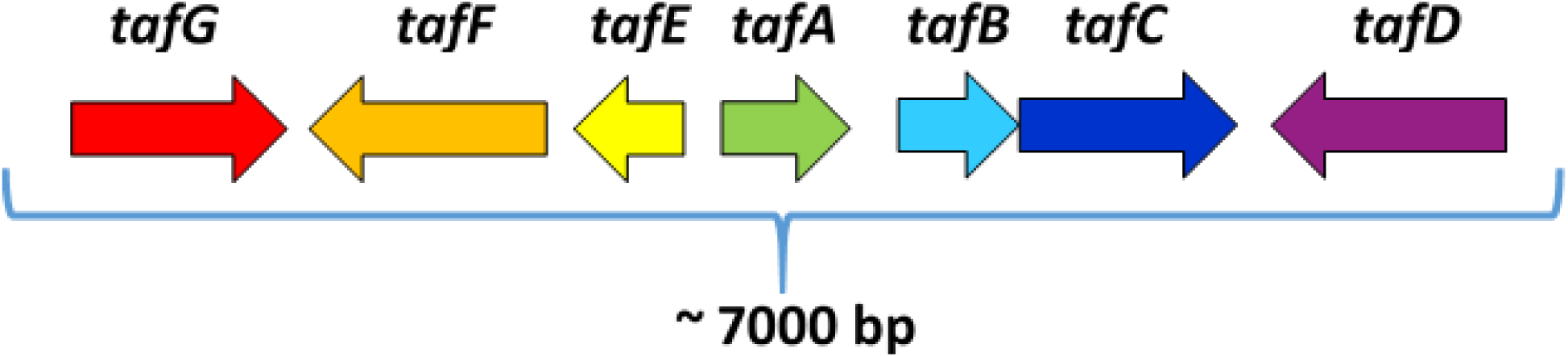
Genetic organization of *Har. hispanica taf* gene cluster *hah_0237-0243* (NC_015948.1 [219076..220188] - NC_015948.1 [complement(224690..225835)]).

If one looks at the annotation of *hah_0239* (*tafE*) and *hah_0242* (*tafC*) genes, their products are hypothetical proteins. TafA and TafE are similar in size and, as it can be seen from a comparison of the predicted secondary structures, are similar structurally (**Figure 2S**). It is possible that TafE performs an adapter function and is responsible for the initiation/termination of fimbria assembly. The N-terminus in TafC domain is similar to TafA and TafE, but its C-terminal part (residues 203-360) includes a laminin-like domain (it refers to concanavalin A-like lectin/glycanases superfamily). It is known that laminins are a large family of adhesive glycoproteins, which are a part of the basal membrane and perform many functions (Domogatskaya *et al*., 2012). Moreover, the proteins of glucanases superfamily participate in binding of sugars and glycoproteins. The presence of such protein in tafi may indicate their adhesive potential and possible involvement in the biofilm formation.

HAH_0241 protein (TafB) is annotated as hypothetical. However, among its close homologues, one, WP_094495324.1 of *Halorubrum ezzemoulense* Ec15, is annotated as Lrp/AsnC family transcriptional regulator. The identity of this protein with TafB is 64%, the positivity is 76%. Lrp/AsnC family of proteins that regulate transcription has been found in both archaea and bacteria. It is known that leucine-responsive regulatory (Lrp) proteins are involved in transport, degradation, and biosynthesis of amino acids; some proteins are involved in the production of pili, porins, sugar transporters, and nucleotide transhydrogenases (Thaw *et al.*, 2006). TafB is predicted to be transmembrane and its functional domain is located in the cytoplasm (**Figure 11**).

**Figure 11.**
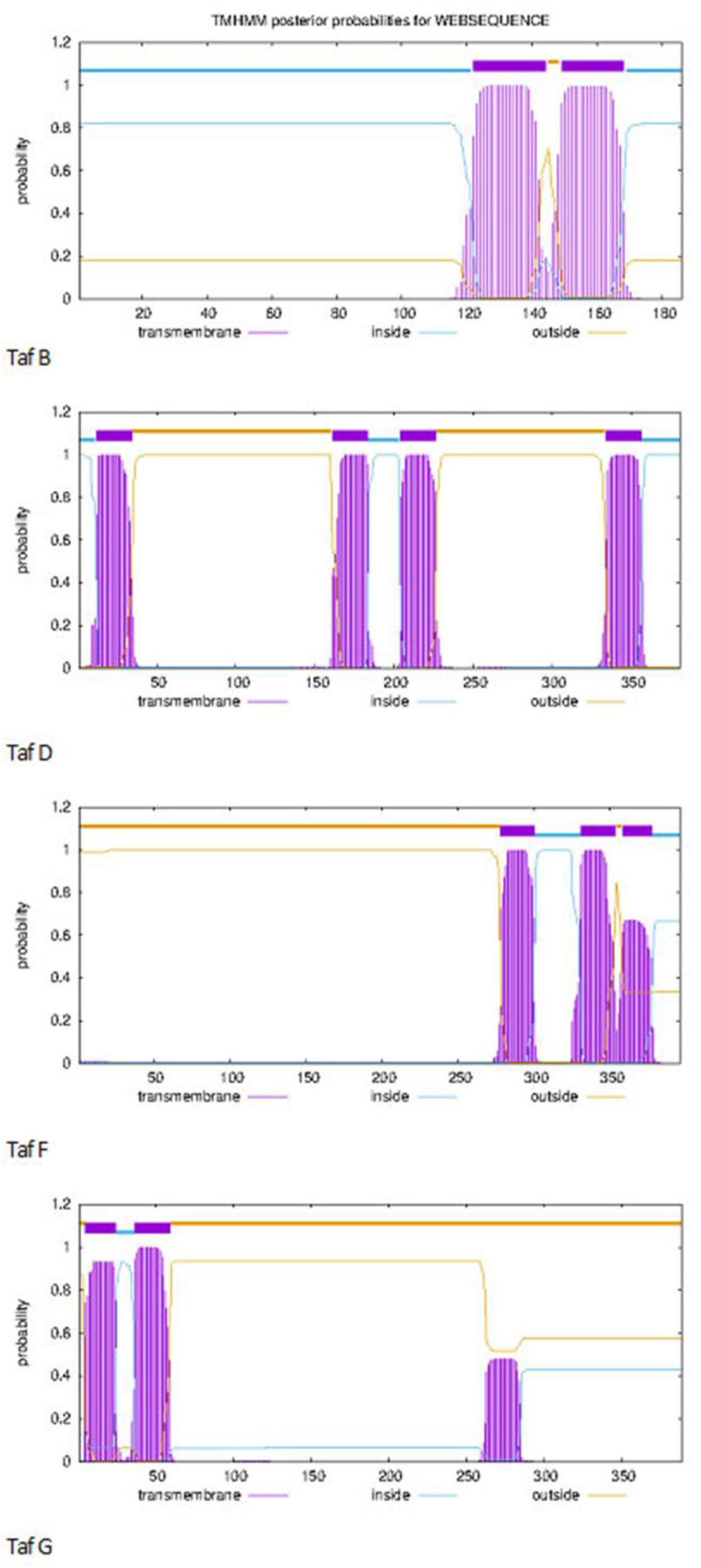
The results of the prediction of transmembrane helices for *Har. hispanica* TafB, D, F, G proteins using TMHMM - 2.0 (Crogh, *et al*., 2001; Hallgren *et al*., 2022).

HAH_0243 (TafD) belongs to S24, S26 superfamily of LexA/signal peptidases, which include membrane-bound proteases that cleave the N-terminal signal peptide of membrane-secreting proteins (Auclair *et al*., 2011). Interestingly, TafD peptidase contains four predicted transmembrane regions and two external domains (**Figure 11**). The sites responsible for peptidase activity are located in the N-terminal domain. The function of the C-terminal domain is unclear. We showed that TafD is involved in the processing of HAH_0240 (TafA) and presumably in the processing of HAH_0239 (TafE) and HAH_0242 (TafC).

HAH_0237 (TafG) contains a DUF5305 domain (amino acid residues 132-349). This DUF5305 family consists of several hypothetical proteins of unknown function and is predominantly represented in archaea and some bacteria (https://www.ncbi.nlm.nih.gov/Structure/lexington/lexington.cgi?cmd=cdd&uid=407348). Interestingly, some bacteria (*Actinoplanes lutulentus* (WP_221402508.1), *Haloplanus salinus* (WP_147270848.1), *Tepidiforma bonchosmolovskayae* (WP_158065712.1), etc.) have signal peptidases, which, along with a domain with peptidase activity, have a DUF5305 domain. In a number of haloarchaea with a taf-cluster, for example, in *Natrinema* sp. SLN56 (**Table 1S**), there are duplicated pairs of genes *tafD* and *tafG*, which may indicate that TafG and signal peptidase TafD function complementary.

HAH_0238 protein (TafF) is annotated as hypothetical. Close homologues of this protein have been found only in archaea with the taf-cluster. One of them, WP_233127167.1 of *Halorubrum* sp. SD612 (**Table 1S**), is annotated as “MSCRAMM family adhesin SdrC”. Its gene is located in the taf-cluster adjacent to genes tafE (WP_086219893.1), tafA (WP_086219895.1) and tafB (WP_086219894.1). The MSCRAMMs (“microbial surface components recognizing adhesive matrix molecules”) are a family of proteins that are defined by the presence of two adjacent IgG-like folded subdomains. These proteins mediate the initial attachment of bacteria to host tissue upon infection. Examples include protein A that binds to IgG, factors from *Staphylococcus* and *Streptococcus* that bind to fibrinogen, etc. (Foster, 2019).

Interestingly, for TafF, the SignalP-5.0 server (Almagro Armenteros *et al*., 2019) predicts with high probability the signal peptide characteristic of the Sec-pathway and not the Tat-pathway, as it does for proteins TafA, TafC, and TafE. For proteins TafB, TafD and TafG, this program does not detect any Tat and Sec signal peptides (**Table 2S**).

Analysis of the protein sequences encoded by the *taf*-gene cluster for the presence of transmembrane regions using the TMHMM-2.0 server (Krogh *et al*., 2001; Hallgren *et al*., 2022; https://services.healthtech.dtu.dk/service.php?TMHMM-2.0) showed that TafB (2 transmembrane helices (TH)), TafD (4 TH), TafF (3 TH) and TafG (3 TH) proteins are probably integrated into the cell membrane (**Figure 11**). Only for TafB, the predominant part of its polypeptide chain is localized inside the cell. For proteins TafD, TafF, and TafG functional domains are predicted to be located on the outer surface of the cell membrane.

Most likely, all of these seven proteins of detected cluster are connected by a common function and participate in the assembly and secretion of tat-fimbriae.

### Molecular modeling of tafi structure (AlphaFold)

By applying AlphaFold Colab (Cramer, 2021; Jumper *et al*., 2021; Mirdita *et al*., 2022) and DALI (Holm, 2020) servers, we predicted and compared the spatial structures of TafA-TafG proteins. It was found that TafA, TafE and N-terminal TafC domain are β-sandwich fold (resembling a canonical jelly-roll fold), most closely related to that of main and auxiliary protein subunits of bacterial adhesive pili I type (P1T). According to analysis using DALI server, TafE shows a greater structural similarity to the main pilus subunit FimA from enteroinvasive bacteria (PDB: 5NKT, 2M5G, 6R74, 6R7E), and TafA to the minor pilin FimG (PDB: 3JWN) and the major pilin subunit CfaB (PDB: 3F83) of enterotoxigenic Escherichia coli. At the same time, the structures of TafA and TafE obtained using AlphaFold fit well into each other (a Z-score = 7.7, %id = 18). Pairwise comparison of TafA/TafE core structures with other Taf-proteins showed the presence of structurally similar domains in N-terminal domain of TafC (for TafA/TafC: Z-score = 3.7, %id = 18), in TafD (for TafA/TafD: Z-score = 6.9, %id = 15) and in TafF (for TafA/TafF: Z-score = 5.8, %id = 9) (**Table 3S, Figures 12, 13**).

**Figure 12.**
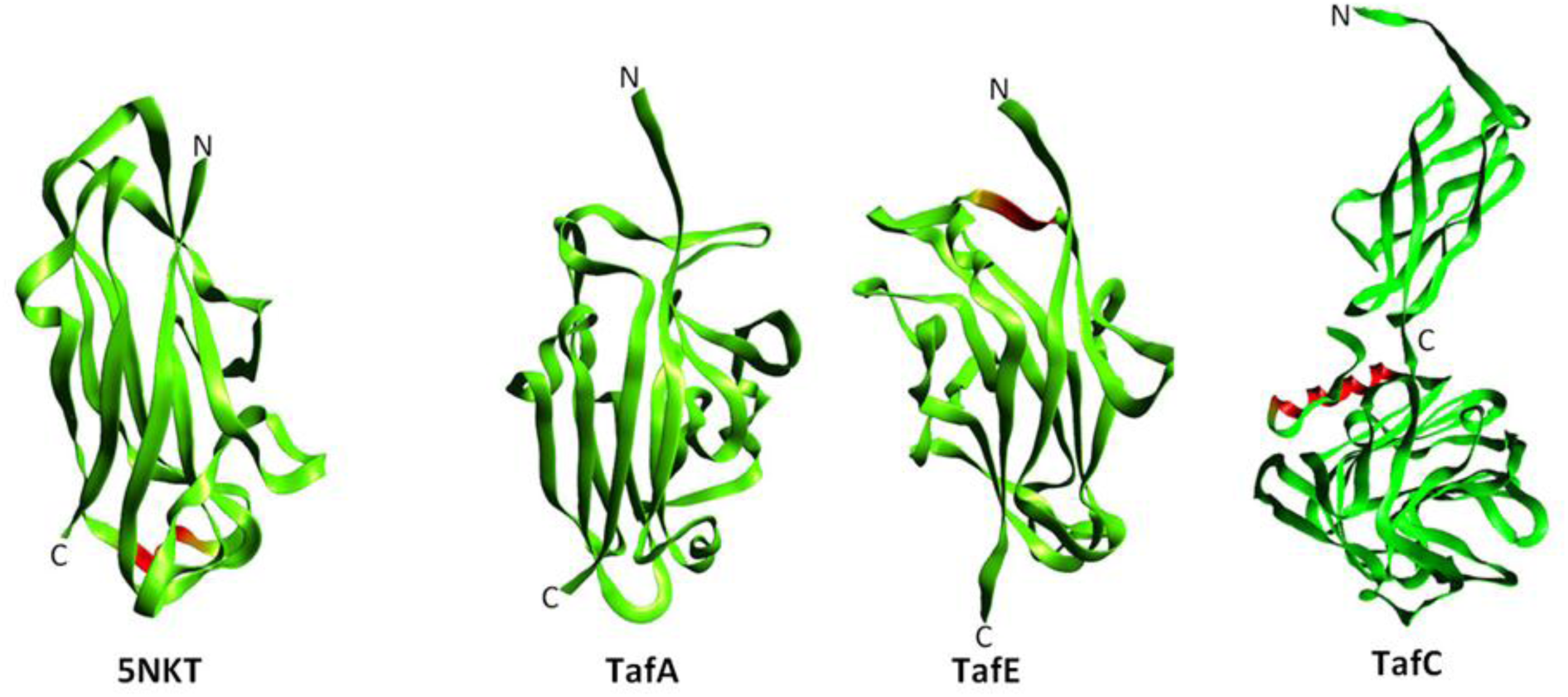
Experimental (X-ray) structure of FimA WT from *E. coli* (5NKT) and Alphafold2 predictions for *Har. hispanica* matured TafA, TafE and TafC proteins.

**Figure 13.**
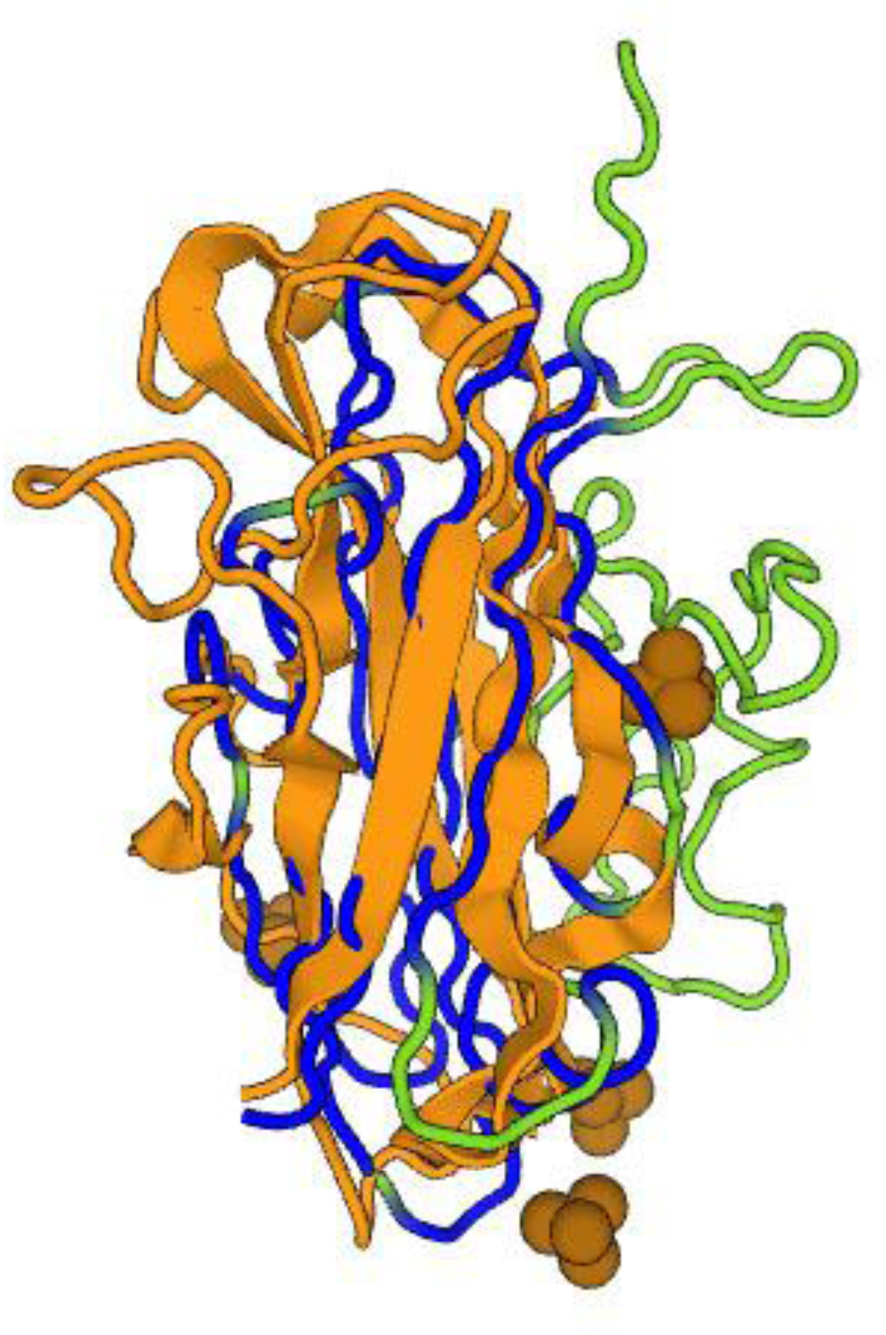
Atomic model of FimA WT from *E. coli* (5NKT) (yellow) superimposed with the Alphafold2 prediction of *Har. hispanica* matured TafA protein (green/blue). TafA regions with a conserved structure are highlighted in blue.

The bacterial adhesive P1T or P-pili are assembled by the chaperone-usher pathway of pilus biogenesis (Lillington *et al*., 2015; Hospenthal *et al*., 2017)). Arrangement of bacterial I type (a) and P pili (b). Adopt from Lillington J. et al. (Biochimica et Biophysica Acta 1840 (2014) 2783-2793). More detailed investigation of P1T composition demonstrated that these structures are multicomponent and consist of several proteins. Individual pilus subunits (FimA/PapA etc.) associate through “donor strand complementation” when the incomplete immunoglobulin-like fold of each subunit is completed by the N-terminal extension of a neighboring subunit. The tip protein (FimH/PapG) is located at the distal pilus end and consists of two crucial domains: the N-terminal lectin domain responsible for adhesive properties and the C-terminal pilin domain structurally similar to the main core protein of pilus (FimA/PapA). Furthermore, there are one or several copies of linker subunits (the proteins FimG, FimF/PapF, PapE, PapK) connecting FimH/PapG with FimA/PapA. About ∼ 1000 copies of FimA/PapA form the rod of pilus. It is important to highlight that all of these proteins use the general Sec-pathway for secretion.

Analyzing our experimental and theoretical data, we suggested that the structural organization of the tafi and adhesive pili is similar, and tafi protomers are also interconnected via donor strand complementation. Indeed, three pivotal protein components (TafA, TafC, TafE) can be thoroughly packed like subunits of the bacterial P1T: FimA, FimH, FimG/FimF, respectively. We have found that TafA is the main component of the filamentous part of tafi. AlphaFold shows that TafC has a two-domain structure: its C-terminal lectin-like domain is probably responsible for the adhesive properties, and the N-terminal domain is involved in polymerization (its structure fits well with that of TafA and TafE).

It is known that there is a special hierarchical arrangement of pilin components in the structure of pilus. As a first step, adhesive proteins FimH/PapG incorporate into pilus and locate on the tip (contact point with the host). Further, the linker protein moves and connects the tip with the rod of pilus. Donor strand complementation is the basic principle underlying mechanism of pilus assembly. This system is fueled by chaperone-usher pathway.

As for tafi, we supposed that they have the same assembly order. On the tip of tafi there is a TafC protein containing a C-terminal adhesive domain (not N-terminal as in pili); then, there is a linker protein TafE, and then, TafA protein that forms the main part of the tat-filament. This assembly order is partly confirmed by our experimental data on the heterologous expression of two genetic constructs: pAS24 with incomplete gene set from *tafA* to *tafD*, and pAS30 with full gene set *tafA* – *tafG*. In the former case, we could not detect any thin filaments, while in the latter case we observed recombinant tat-fimbria. Does it mean that TafA and TafC cannot connect directly with each other and assemble the filament and need an intermediary protein (TafE) between them? One more confirmation of our suggestion is the deletion of tafE gene that leads to disturbance of TafA protein synthesis (there is no TafA protein band on SDS-electropherogram) and presumably to termination of tafi assembly (data not shown).

Using the AlphaFold server, we obtained hypothetical structures of oligomers of subunits TafA, TafC and TafE. Interestingly, when considering homodimers of the TafA or TafE subunits, AlphaFold does not predict interactions via donor strand complementation. However, in the model of the structure obtained for the heteromer TafC/TafE, an interaction via complementation of the TafC N-terminal donor chain with TafE with outward protrusion of the TafE donor chain takes place. However, in the model of the interaction of TafC/TafA, the protrusion of the TafA donor chain does not occur (**Figure 14**). Modeling for tetramers TafC/2xTafA/TafE and TafC/TafA/2xTafE demonstrated filamentous complexes of linearly interacting molecules: in this case, TafC is located at the end; its donor chain interacts with TafE, activating it; and TafE, in turn, interacts with molecules of TafE or TafA via its donor chain. The main subunit TafA has a lower affinity for TafC than protein TafE and requires TafE presence to activate its ability to polymerize.

**Figure 14.**
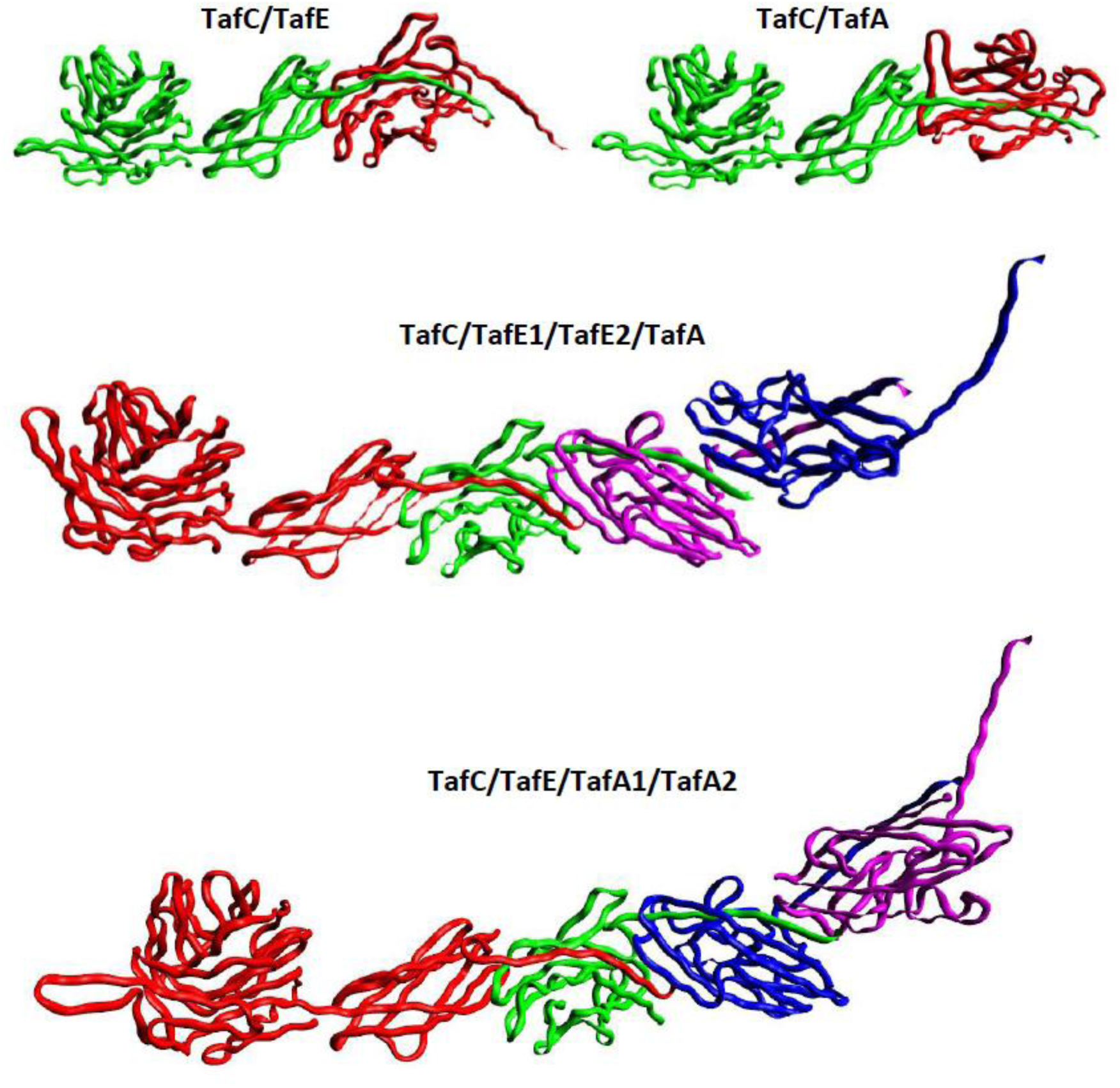
Alphafold2 predictions for heteromeric complexes of *Har. hispanica* matured TafA, TafE and TafC proteins.

**Figure 15** shows the model for the TafC/TafE/TafA/TafF heterotetramer made with AlphaFold. As we have already mentioned, the N-terminal domain of TafF is structurally similar to the domains of key subunits TafA, TafC and TafE. It can be seen that the interaction of the complementary donor chain of TafA with the N-terminal TafF domain can take place. Since TafF contains C-terminal transmembrane regions, it can be assumed that it is responsible for tafi anchoring in the cell membrane.

**Figure 15.**
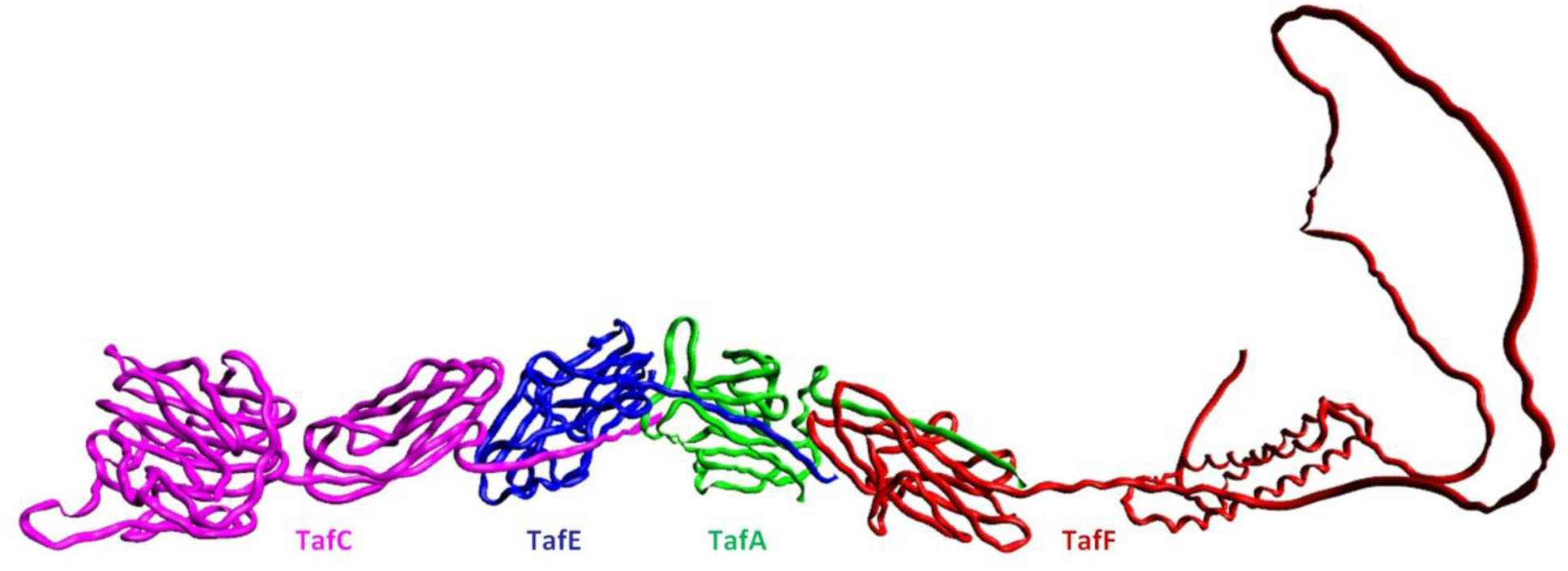
Alphafold2 predictions for heterotetrameric complexes of *Har. hispanica* matured TafA, TafC, TafE and TafFproteins.

The AlphaFold reveals that the most appropriate model for tat-fimbria is the structure from three proteins arranged as follows: TafC on the tip, TafE –as linker, and TafA as the core of filament (**Figures 14, 15**). However, we do not know yet how TafA anchor to the cell wall and how the filament assembly is regulated. Are there some proteins that can play the role of usher or chaperone? We presuppose that other proteins from the TafA-TafG cluster can be similar to these and also participate in this process. Future studies should shed light on these questions.

## CONCLUSION

Although an increasing number of archaeal surface appendages are being identified, the function of only a few ones has been studied in detail. Among these structures are Iho670 fibers, Mth60 fimbriae, hami, cannulae, bindosomes, etc. (Jarrell and Albers, 2012; Jarrell *et al*., 2013). Usually, these structures were found by using electron microscope observations. It is often very difficult to describe their functions and structural features due to the complexity of strain cultivation and lack of data on protein composition.

The development of the unique method of cryo-electron microscopy and new bioinformatic servers such as AlphaFold and DALI have remarkably improved the possibilities of studying the structure and mechanisms of various biological supramolecular complexes. Recently, two teams of researchers reported on the atomic structures of two previously uncharacterized archaeal surface filaments: the bundling pili of *Pyrobaculum calidifontis* (Wang *et al*., 2022) and the thread filament of *Sulfolobus acidocaldarius* (Gaines *et al*., 2022) determined using the cryo-electron microscopy and AlphaFold. Both filamentous structures show a striking resemblance to type I bacterial adhesive pili.

In this study, we reported on the discovery and primary analysis of the novel type of haloarchaeal surface structures. We named them “tat-fimbriae – tafi”, thereby emphasizing the uniqueness of the secretion pathway of the main structural components that these filaments are composed. We found that the pivotal structural preproteins of tafi – TafA, TafC, TafE – contain a specific twin-arginine motif that is usually detected in plant, bacterial and archaeal proteins that use Tat-pathway for secretion. All currently known surface structures of archaea use the general secretion way (Sec-pathway).

We speculate that the use of the Tat-pathway for the assembly of surface filamentous structures in haloarchaea is quite expected. It is known that in haloarchaea many surface proteins (various extracellular enzymes, proteins responsible for adhesion and redox processes, etc.) most often use Tat-pathway for secretion. It is a favored way of evolutionary adaptation to highly saline conditions.

Structural similarity of the Taf proteins to the proteins of type I bacterial pili is drawing particular interest. Despite the differences in the amino acid sequence of the main proteins of tafi and pili (and despite the lack of homology between proteins), the principle of formation (a common subunit packing) of such thin filamentous surface structures is very similar in different species from Bacteria and Archaea domains.

## EXPERIMENTAL PROCEDURES

### Strains and growth conditions

The plasmids and strains used in this study are listed in **Table 1**.

**Table 1:** Plasmids and strains.

| Plasmid/Strain | Relevant properties | Reference |
| --- | --- | --- |
| Plasmids |  |  |
| pHAR | 4.0 kb; vector for gene knockout $Amp^r$, *pyrF<sub>Hh</sub>* and its native promoter | Liu *et al.*, 2011a |
| pAG1 | vector based on pHAR for deletion *tafA* (*hah_0240*) | This study |
| pAG2 | vector based on pHAR for deletion *tafD* (*hah_0243*) | This study |
| pAG3 | vector based on pHAR for deletion *arlK* (*hah_2955*) | This study |
| pTA1228 | inducible vector for *Hfx. volcanii*, $Amp^r$, *pyrE2* and *hdrB* markers, inducible *ptna* promoter | Haque *et al.*, 2020 |
| pAS24 | vector based on pTA1228 with gene cassette *tafA- tafD* from *Har. hispanica*. *tafA* gene is located under *ptna* promoter, the other genes are under their own promoters | This study |
| pAS30 | vector based on pAS24, with full gene cassette *tafA- tafG* | This study |
| pAS31 | pAS30 containing *tafA* with R5K and R6K mutations | This study |
| Strains |  |  |
| DH5α | *E. coli* F<sup>-</sup> ϕ80*lacZDM15* (*lacZYA-argF*) U169 *recA1 endA1 hsdR17* (*rK<sup>-</sup> mK<sup>+</sup>*) *phoA supE44 thi-1 gyrA96 relA1* | Invitrogen |
| XL1-Blue | *E. coli* *recA1 endA1 gyrA96 thi-1 hsdR17 supE44 relA1 lac* [*F'**proAB lacI<sup>q</sup>ZΔM15 Tn10* (*Tet<sup>r</sup>*)] | Evrogen |
| WT | Wild-type *Haloarcula hispanica* strain (B- | Juez *et al.*, 1986; Liu *et al.*, |
|  | 1755, DSM 4426, ATCC 339606 CGMCC 1.2049) | 2011b |
| DF60 | *Har. hispanica* ΔpyrF | Liu *et al.*, 2011a |
| AG | DF60 $\Delta$ *tafA* (*hah_0240*) | This study |
| AG1 | DF60 $\Delta$ *tafD* (*hah_0243*) | This study |
| AG2 | DF60 Δ*arlK* | This study |
| AG3 | AG2 Δ*tafA* | This study |
| AG4 | AG2 Δ*tafD* | This study |
| MT78 | *Hfx. volcanii* ΔpyrE2 Δtrp ΔallA1ΔarlA2 ΔpilB3-C3 | Esquivel *et al.*, 2014 |
| AS24 | MT78 containing pAS24 | This study |
| AS30 | MT78 containing pAS30 | This study |
| AS31 | MT78 containing pAS31 | This study |

*Haloarcula hispanica* strain B-1755 (DSM 4426, ATCC 33960, CGMCC 1.2049) was obtained from the All-Russian Collection of Microorganisms, (VKM) Pushchino. We marked it as a wild type (WT) strain. The *Har. hispanica* strain DF60 was constructed from the WT strain by deletion of *pyrF* gene (encoding the orotidine – 5’ phosphate decarboxylase) as described by Liu *et al*. (2011a). *Har. hispanica* strains were cultivated at 37°С or 42°С in a complex medium AS-168 (per 1L: 5 g Bacto Casamino acids, 5 g yeast extract, 1 g sodium glutamate, 3 g trisodium citrate, 20 g MgSO_4_*7H_2_O, 2 g KCl, 200 g NaCl, 50 mg FeSO_4_*7H_2_O, 0.36 mg MnCl_2_*4H_2_O, pH 7.0) (Han *et al.*, 2007). We also used the AS-168SY medium similar to the AS-168, except that yeast extract was omitted. When required, AS-168SY medium was supplemented with uracil (Sigma-Aldrich) at a concentration of 50 μg/ml, and 5-FOA (Thermo Fisher Scientific) at 150 μg/ml.

*Haloferax volcanii* strain MT78 (Esquivel and Pohlschröder, 2014) was kindly provided by M. Pohlschröder. All *Hfx. volcanii* transformed strains were grown at 37°C in liquid or agar (0.5% for solid and 0.24% for semi-solid) medium (per 1L: 5 g Bacto Casamino acids, 200 NaCl, 18 g MgSO_4_, 4 g KCl, 0.33 g CaCl_2_, 0.2 g MnCl_2_*4H_2_O, pH 7.2). Heterologous *tafA*-gene expression was induced by adding tryptophan (0.2-1 mg/ml). The rest of the taf-genes were expressed under their natural promoters.

*Escherichia coli* XL1-Blue was grown in LB medium with added 100 μg/ml of ampicillin.

### Isolation and separation of archaella and tafi

Normally, for isolation of archaellar filaments we used polyethylene glycol precipitation (PEG 6000) as described by Gerl *et al*. (1989). However, we found that *Har. hispanica* archaella and tafi become co-isolated if this procedure is used. Differential centrifugation was used to separate the two types of filaments. At first, we pelleted the cells by low-speed centrifugation at 8000 g using Beckman J2-HS centrifuge and the JA-14 fixed angle rotor. The resulting supernatant contained discarded archaella and tafi. Archaella were precipitated from the supernatant at 142400 g using Beckman Coulter Opima XE-90 ultacentrifuge and the fixed-angle titanium rotor Type 45 Ti (Beckman Coulter). The tafi remained in the supernatant and could be precipitated either during many hours of high-speed centrifugation, or within 1 hour at 46500 g after adding 4% polyethylene glycol 6000. The resulting preparation contained a relatively pure tafi; the yield was approximately 5 mg/1 liter of liquid culture. Precipitated archaella and tafi were dissolved in 10 mM Tris–HCl, pH 8.0, containing 20% NaCl. SDS-PAGE (sodium dodecyl sulfate–polyacrylamide gel electrophoresis) was performed using 12% acrylamide gels. The proteins were stained with Coomassie Brilliant Blue G-250.

A similar procedure using PEG precipitation was used to isolate recombinant tafi from *Hfx. volcanii* MT78. Although this strain does not synthesize its own archaella and pili, preliminary high-speed centrifugation was useful to remove fragments of cell membranes and other contaminating material. When the plasmid pAS24 was expressed in *Hfx. volcanii* MT78, the cells were peletted by low-speed centrifugation at 8000 g using Beckman J2-HS and the supernatant was additionally centrifuged at 142400 g using Beckman Coulter Opima XE-90 ultacentrifuge (twice) to remove contaminating material. Then it was concentrated using Amicon Ultra Centrifugal Filters Ultracel – 3K (Merck) according to the manufacturer’s instructions. The original volume (50 ml) was concentrated to a volume of approximately 500 μl (100 times). The presence of target proteins in the extracellular medium was detected using SDS-electrophoresis and mass spectrometry.

### Mass spectrometry analysis

Mass spectra of the samples were obtained using an OrbiTrap Elite mass spectrometer (Thermo Scientific, Germany). The procedure for sample preparation, measurements, and data analysis was described previously (Pyatibratov *et al*., 2020).

### Electron microscopy

Archaella and tafi samples were prepared for transmission electron microscopy (EM) by negative staining with 2% uranyl acetate on Formvar-coated copper grids. A grid was floated on a 20-μl drop of archaella or tafi solutions (about 0.01-0.05 mg/ml, in 20% (3.4 M) NaCl, 10 mM TrisHCl, pH 8.0) for 2 min, blotted with filter paper and placed on top of a drop of 2% uranyl acetate for 1 min. Excess stain was removed by touching the grid with the filter paper, and the grid was then air dried. Samples were examined on a Jeol JEM-1400 transmission electron microscope (JEOL, Japan) operated at 120 kV. Images were recorded digitally using a high-resolution water-cooled bottom-mounted CCD camera.

### TafA (HAH_0240) purification

Additional purification of TafA (HAH_0240) protein from archaella traces, high-molecular membrane proteins, and other impurities was carried out by depolymerization of Tat strands followed by size-exclusion chromatography on a Superose 12 column. For this, after PEG precipitation tafi preparation was diluted in 50 mM Tris-HCl buffer with 8.0 pH containing 0-5% NaCl and kept for 15 minutes in a thermostat at 90°C. Then, the preparation was cooled and centrifuged for 30 min at 213400 g in a TLA-100 centrifuge, Beckman. The supernatant was applied to a Superose 12 column equilibrated with 10 mM Na-phosphate buffer containing 0, 5, 10, 20, or 25% NaCl. The elution rate at all stages of the experiment was 0.2 ml/min. Protein molecular weights were estimated from elution volumes using BSA (66 kDa) (Sigma-Aldrich) and ovalbumin (45 kDa) (Sigma-Aldrich) as standards. The presence of proteins at the outlet of the column was recorded using spectrometry followed by SDS electrophoresis. The vast majority of contaminant proteins, including archaellins, elute directly behind the column void volume, while TafA protein elutes with a significant delay. The study of the secondary structure of TafA was carried out using circular dichroism (CD) spectroscopy on a Chirascan spectropolarimeter (Applied Photophysics) in the wavelength range from 190 nm to 250 nm. The values of molar ellipticity were calculated following the equation: [θ] = [θ]_m_ М_res_/(LC), in which C is the protein concentration (mg/ml), L is the optical path length of the cuvette (mm), [θ]_m_ is the measured ellipticity (degrees) and М_res_ is the average molecular weight of the peptide residue (Da) calculated from its amino acid sequence. Measurements were carried out in a 0.1 mm cuvette. Calorimetric measurements were carried out on a SKAL-1 differential scanning microcalorimeter (ZAO SKAL, Pushchino, Russia) at a heating rate of 1°C/min. Working volume of the gold cell was 0.3 ml; protein concentration in the samples used in the experiments was 1–0.5 mg/ml. For the experiments, we used 10 mM Na-phosphate buffer, pH 8.0, containing 25% NaCl. Measurements were carried according to the method described earlier (Privalov and Potekhin, 1986; Tarasov *et al*., 1995).

### Construction and confirmation of the deletion mutants

Chromosomal deletions in AG, AG1 and AG2 strains were generated using homologous recombination (pop-in/pop-out) method as previously described (Liu *et al*., 2011a). The sequences of PCR primers used in this study were summarized in Table 2.

**Table 2:**
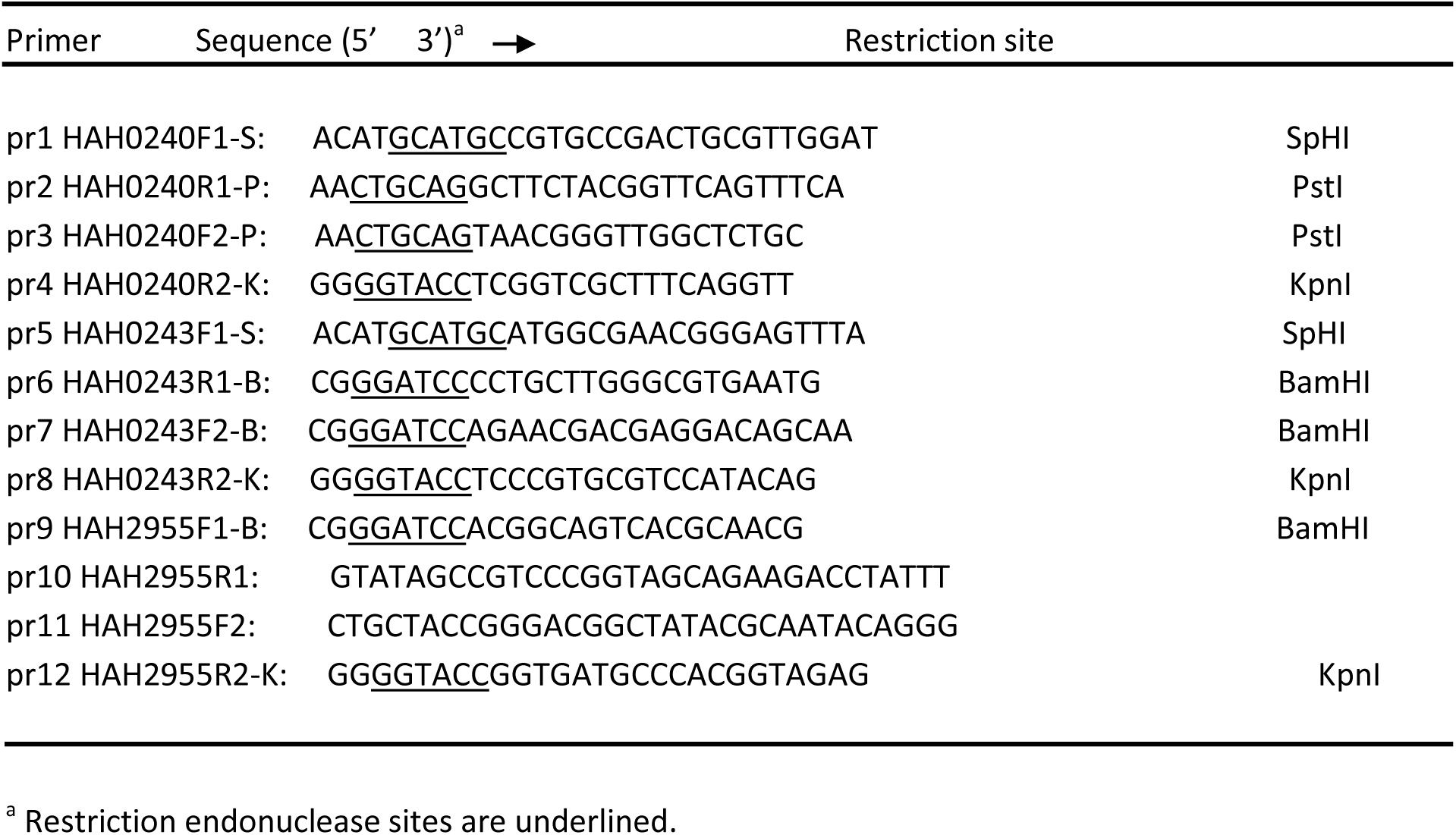
Oligonucleotides used in this study.

To obtain the target strains, we prepared a pair of DNA fragments by PCR using primers pr1/pr2 and pr3/pr4 for AG strain; pr5/pr6 and pr7/pr8 for AG1 strain; pr9/pr10 and pr11/pr12 for AG2 strain. In the case with AG and AG1 strains, two amplified DNA fragments (up- and downstream flanking sequences for each gene) were cut with corresponding restriction endonucleases and mixed together with the pHAR plasmid treated by the same restrictases. After ligation and transformation, the colonies of E. coli XL1-Blue competent cells were screened by PCR and then those containing pHARΔ*tafA*/pHARΔ*tafD* were selected.

For AG2 strain, two amplified DNA fragments were linked using overlap PCR with a pair of primers of pr9 and pr12 to generate a fragment containing corresponding sites for restriction (Table 2). The fragment was then cloned into the cut pHAR plasmid, too. Out of the competent cells, those with the pHARΔ*arlK* plasmid were selected.

The next step was *Har. hispanica* DF60 transformation with plasmids pHARΔ*tafA*/ pHARΔ*tafD*/ pHARΔ*arlK*. Cells were plated onto AS-168SY solid medium (a nutrient-poor medium without yeast extract). Transformants were screened for integration of the gene knockout plasmid at the corresponding locus by PCR analysis. Cells with required plasmids pHAR(Δ*tafA*/Δ*tafD*/ or Δ*arlK*) integrated into their genome were subcultured into AS-168 medium (complex medium with yeast extract) supplemented with 50 μg/ml Uracil and 150 μg/ml 5-FOA for counterselection of the double-crossover recombinants.

PCR products were analyzed by agarose gel electrophoresis.

### Heterologous expression of Har. hispanica tafA-tafG genes in Haloferax volcanii

For heterologous expression, we used the special strain of *Hfx. volcanii* MT78, in which the archaellin operon (Δ*arlA1*Δ*arlA2*) and genes critical for the synthesis of type IV pili (*ΔpilB3-C3*) were removed (Esquivel *et al*., 2014). For inducible expression, vectors based on pTA1228 with tryptophan inducible promoter (Allers *et al*., 2010) were used.

To obtain the expression plasmid pAS24, a fragment containing *tafA, B, C, D* genes was generated using PCR. Further, the original pTA1228 vector was cleaved with EcoRI and NdeI enzymes and, using the TEDA method, (Xia *et al*., 2019) the resulting PCR fragment was inserted into it. To obtain the pAS30 vector, the pAS24 plasmid was cleaved at the NotI site, and then, using the TEDA method, generated PCR fragment containing *tafG, F*, and *E* genes was inserted into it.

To obtain the pAS31 vector, pAS30 plasmid was cleaved at NdeI and KasI sites, followed by insertion of the PCR fragment generated with primers pAS31_F1 and pAS31_R1.

The correctness of the assembly of all obtained plasmids was checked using Sanger sequencing.

### Structural prediction (Alphafold2)

Alphafold2 (Cramer, 2021; Jumper *et al*., 2021) was used to predict the structures of Taf-proteins (HAH_0237-HAH_0243), using the online ColabFold (Mirdita *et al*., 2022) tool. Structure alignment was performed by the Dali server (Holm, 2020). Protein model structures were viewed using Avogadro tool (Hanwell *et al*., 2012; http://avogadro.cc/).

## Acknowledgements

This work was supported by a RFBR grant No. 19-04-01327 A to A.S. We thank Dr. T. Allers for pTA1228 plasmid and Dr. M. Pohlschröder for the kindly presented strains and M. Suvorina for the help with MS analysis. The authors thank B.S. Melnik for CD measurements and T.N. Melnik for calorimetric measurements. Electron microscopy was performed with the support of Moscow State University development program (PNR 5.13). Mass spectrometric analysis was performed in facilities of United Pushchino Center “Structural and functional studies of proteins and RNA” (584307).

## Author contributions

A.V.G., M.G.P., A.S.S. and H.X. conceived of the project, designed the study and wrote the paper; A.V.G., A.S.S., M.G.P., D.Z., E.Yu.P., J.L., I.I.K. and A.K.S. performed the experiments; A.V.G., M.G.P., A.S.S. and A.K.S. analyzed the data; A.S.S., H.X., M.G.P., I.I.K. and A.K.S. contributed funding and resources.

## Conflict of interest

The authors declare that they have no conflict of interests.

## TABLES AND FIGURE LEGENDS

**Table 1S.** A list of close homologues of Har. hispanica ATCC 33960 TafA (HAH_0240) found using the BLAST program. The proteins are ranked in decreasing order of their homology with HAH_0240. For each of the organisms, the found homologues of the tafA-G cluster genes are indicated. Various paralogs of the corresponding genes are numbered. Species in which all 7 genes of the taf-A-G cluster were found are in bold.

**Table 2S.** Signal peptide and cleavage sites prediction results for TafA-TafG proteins using the SignalP-5.0 server (Almagro Armenteros *et al*., 2019).

**Table 3S.** Results of pairwise comparison of Har. hispanica TafA-TafG protein structures predicted using AlphaFold and 5NKT, X-ray structure of FimA wt from E. coli (Zyla et al., 2019) using DALI server. Z-score (similarities with a Z-score lower than 2 are spurious) and percent identity (%id).

**Figure 1S.** Mass spectrometry analysis of major protein band in tafi preparation isolated from *Har. hispanica* DF60. Protein coverage (86%) of HAH_0240 (WP_044951594.1) is shown. In gray the recognized protein sequence is shown. In blue the unique peptides are depicted.

**Figure 2S.** Prediction of the secondary structures of proteins TafA using PSIPRED protein structure prediction server http://bioinf.cs.ucl.ac.uk/psipred/ (Buchan *et al*., 2019).

**Figure 3S.** The pAS24 plasmid construction with *tafA-tafD* genes from *Har. hispanica* for heterologous expression in *Hfx. volcanii*.

**Figure 4S.** Mass spectrometry analysis of protein bands 1-4 in preparation obtained by expression of pAS24 plasmid containing *Har. hispanica tafA, tafB, tafC* and *tafD* genes in a heterologous host *Hfx. volcanii*. *Har. hispanica* TafA (HAH_0240, WP_044951594.1), TafC (HAH_0242, WP_014039247.1) proteins and *Hfx. volcanii* HVO_A0133 protein were detected in all bands 1-4. The protein coverages of TafA (60%, 82%, 87% and 79%, respectively), TafC (55%, 15%, 12% and 26%, respectively) and HVO_A0133 (18%, 52%, 73% and 83%, respectively) are shown. In gray the recognized protein sequence is shown. In blue the unique peptides are depicted.

**Figure 5S.** The pAS30 plasmid construction with *tafA-tafG* genes from *Har. hispanica* for heterologous expression in *Hfx. volcanii*.

**Figure 6S.** Conserved tafA-neighbouring gene clusters in haloarchaea with fully sequenced genomes. Similar genes are painted in the same color: *tafA* - green, *tafB* - blue, *tafC* - dark blue, *tafD* – purple, *tafE* – yellow, *tafF* - orange, *tafG* - red. The *Hfx. mediterranei* genome contains a cluster with the same arrangement of taf-genes as in Har. hispanica. Har. marismortui genome contains 2 taf-clasters: a complete set of 7 *tafA-tafG* genes (+ an additional *tafG* gene) on chromosome 1, and an incomplete set (genes *tafA, tafC, tafE* and *tatF*) on plasmid pNG200. The picture was created using the Gene Graphics server (Harrison *et al*., 2018).

**Figure 7S.** Mass spectrometry analysis of minor protein bands in tafi preparation isolated from *Har. hispanica* DF60. (A) - the result of identification for the band above protein HAH_0240, (B) - the result for the band below HAH_0240. Protein coverage of HAH_0242 (WP_014039247.1) and HAH_0239 (WP_044951591.1) are shown (79% and 61%, respectively). In gray the recognized protein sequence is shown. In blue the unique peptides are depicted. The sequence from the database used for protein identification contains a truncated version of protein HAH_0239. A more likely variant is 14 amino acid residues longer, starts with a previous start codon and contains a twin arginine pattern.

## Notes

### Competing Interest Statement

The authors have declared no competing interest.

