## Supplementary for "Tat-fimbriae (“tafi”) – novel type of haloarchaeal surface structures"

| | Archaea | Sequence ID | Identities, % | Positivies, % | Cluster of taf-genes |
|---|---|---|---|---|---|
| 1 | **Haloarcula hispanica ATCC 33960** | WP_014039245.1, AEM55868.1 | 100 | 100 | tafA,B,C,D,E,F,G |
| 2 | **Haloarcula hispanica N601** | WP_014039245.1, AHB64694.1 | 100 | 100 | tafA,B,C,D,E,F,G |
| 3 | **Haloarcula hispanica CBA1121** | KAA9408629.1 | 98 | 98 | tafA,B,C,D,E,F,G |
| 4 | **Haloarcula sp. K1** | KZX49737.1 | 97 | 98 | tafA,B,C,D,E,F,G |
| 5 | **Haloarcula sp. CBA1122** | MUV50900.1 | 97 | 98 | tafA,B,C,D,E,F,G |
| 6 | **Haloarcula hispanica 18.1** | RYJ10990.1 | 97 | 98 | tafA,B,C,D,E,F,G |
| 7 | **Haloarcula sp. CBA1115** | AJF25871.1 | 98 | 98 | tafA,B,C,D,E,F,G |
| 8 | **Haloarcula sp. CBA1128** | WP_175055151.1 | 98 | 98 | tafA,B,C,D,E,F,G |
| 9 | **Haloarcula sp. CBA1131** | KAA9405491.1 | 91 | 93 | tafA,B,C,D,E,F,G |
| 10 | **Haloarcula sp. Atlit-7R** | WP_195156628.1, RLM88247.1 | 89 | 92 | tafA,B,C,D1,E,F,G1 + tafD2G2 + tafD3G3 |
| 11 | **Haloarcula sp. Atlit-120R** | RLM36418.1 | 89 | 92 | tafA,B,C,D,E,F,G |
| 12 | **Haloarcula sp. Atlit-47R** | RLM45200.1 | 88 | 92 | tafA,B,C,D1,E,F,G1 + tafD2G2 |
| 13 | **Haloarcula taiwanensis** | AUG46196.1 | 88 | 92 | tafA,B,C,D1,E,F,G1 + tafD2G2 |
| 14 | **Haloarcula sp. CBA1127** | WP_195156628.1 | 83 | 89 | tafA,B,C,D,E,F,G |
| 15 | **Haloarcula amylolytica JCM 13557** | WP_008311393.1, EMA19641.1 | 82 | 87 | tafA,B,C,D,E,F,G |
| 16 | **Haloarcula rubripromontorii pws8** | WP_170084197.1, NLV07818.1 | 82 | 90 | tafA,B,C,D1,E,F,G1 + tafD2G2 |
| 17 | **Haloarcula rubripromontorii SL3** | WP_053966619.1, KOX94854.1 | 82 | 90 | tafA,B,C,D,E,F,G |
| 18 | **Haloferax larsenii JCM 13917** | WP_007542347.1, ELZ79117.1 | 69 | 81 | tafA,B,C,D,E,F,G |
| 19 | **Haloferax elongans ATCC BAA-1513** | WP_008327219.1, ELZ80264.1 | 68 | 81 | tafA,B,C,D,E,F,G |
| 20 | **Haloferax larsenii CDM_5** | WP_074795815.1, SEL81368.1 | 68 | 81 | tafA,B,C,D,E,F,G |
| 21 | **Haloferax mediterranei ATCC 33500** | WP_004061029.1, AFK21216.1 | 67 | 79 | tafA,B,C,D,E,F,G |
| 22 | Halorubrum sp. SD683 | WP_086216962.1, OTE98601.1 | 41 | 57 | no data |
| 23 | Halorubrum ezzemoulense Ga2p | WP_094593028.1, OYR67436.1 | 42 | 57 | tafA,B,C,D1,F + distant cluster containing tafD2,G. tafE? |
| 24 | Halorubrum sp. ARQ200 | WP_137704375.1, TKX45036.1 - **tafA1** | 42 | 55 | tafA1,C,D1,F1 + distant tafB,D2,G + tafA2,D3,E,F2 |
| 25 | Halorubrum sp. ASP121 | WP_137704375.1, TKX48818.1 | 42 | 55 | tafA,C,D1,F + tafB,D2,G |
| 26 | **Halorubrum sp. PV6** | WP_168654444.1, AZQ16086.1 | 43 | 55 | tafA,B,C,D1,E,F + tafD2,G1 + tafD3,G2 + tafD4,G3 |
| 27 | Halorubrum sp. SD612 | WP_086219894.1, OTF10344.1 | 42 | 57 | tafA,B,C,D1,E,F + tafD2,G |
| 28 | Halorubrum sp. GN12_10-3_MGM | WP_137708496.1, TKX62663.1 | 43 | 57 | tafA,B,C,D1,E,F + tafD2,G |
| 29 | Halorubrum litoreum JCM 13561 | WP_008368288.1, EMA57595.1 | 37 | 54 | tafA,B1,C,D1,F + distant tafD2,G + tafB2,E,D3,G |
| 30 | Halorubrum terrestre 22517_05_Cabo | WP_159369376.1, MYL17491.1 - **tafA1** | 39 | 53 | tafA1,A2,B1,C,D1,E,F + tafB2,D2,F2 + tafD3,G |
| 31 | **Halorhabdus sp. CBA1104** | WP_154552456.1, QGN07449.1 | 43 | 57 | tafA,B,C,D,E,F,G |
| 32 | **Halorubrum sp. WN019** | WP_095638187.1, PAU80007.1 | 40 | 51 | tafA,B,C,D1,E,F + tafD2,G |
| 33 | Halorubrum coriense DSM 10284 | WP_049910780.1 | 40 | 54 | tafA,B,D1,E,F + tafD2,G |
| 34 | Halorubrum ezzemoulense Ec15 | WP_094495325.1, OYR72685.1 | 36 | 52 | tafA,B,C,D1,F + tafD2,G |
| 35 | Halorubrum sp. Ea1 | WP_094558349.1, OYR50889.1 | 40 | 54 | tafA,B,C,D1,F + tafD2,G,E |
| 36 | Halorubrum sp. Ib24 | WP_094581915.1, OYR38331.1 | 39 | 53 | tafA,B,C,D,F |
| 37 | Haloarcula vallismortis DSM 3756 | WP_049919878.1, SDW00890.1 | 41 | 56 | tafA,B,C,D,E,F,G |
| 38 | Halorubrum sp. Hd13 | WP_094590236.1, OYR40738.1 | 37 | 52 | tafA1,B,C,D1,F + tafA2?,D2,G |
| 39 | Halorubrum tebenquichense DSM 14210 | WP_082238212.1 | 40 | 56 | tafA,B,C,D1,E,F + distant cluster containing tafD2,G genes |
| 40 | Halorubrum sp. Ea8 | WP_094527960.1, OYR45145.1 | 37 | 53 | tafA,B,C,D1,?,? + tafD2,G |
| 41 | Halorubrum ezzemoulense Ga36 | WP_143420758.1, OYR61783.1 | 35 | 51 | tafA,B,C,D1,F + tafD2,G |
| 42 | Halorubrum sp. Eb13 | WP_094523749.1, OYR45996.1 | 38 | 54 | tafA,B,?,D1,?,? + tafD2,G |
| 43 | Halorubrum chaoviator DSM 19316 | SNR61959.1 | 37 | 50 | tafA,B,C,D1,F + tafD2,G |
| 44 | Halorubrum sp. CGM4_25_10-8A | WP_137703867.1, TKX36926.1 | 37 | 50 | tafA,B,C,D1,F + tafD2,G |
| 45 | Halorubrum ezzemoulense DSM 17463 | WP_139835076.1 | 36 | 53 | tafA,B,C,D1,F + tafD2,G1 + tafD3,G2 |
| 46 | **Halorubrum sp. 48-1-W** | WP_112080037.1, RAW44336.1 | 38 | 51 | tafA,B,C,D1,E,F + tafD2,G |
| 47 | **Haloarcula sp. CBA1129** | WP_151101155.1, KAA9399212.1 | 39 | 54 | tafA,B,C,D,E,F,G |
| 48 | **Haloarcula sp. CBA1130** | WP_151101155.1, KAA9403726.1 | 39 | 54 | tafA,B,C,D,E,F,G |
| 49 | **Halorubrum hochstenium ATCC 700873** | WP_008585674.1, ELZ54024.1 | 45 | 56 | tafA,B,C,D1,E,F + tafD2,G |
| 50 | **Haloarcula argentinensis pws5** | WP_170097738.1, NLV14318.1 | 39 | 54 | tafA,B,C,D,E,F,G |
| 51 | **Haloarcula argentinensis DSM 12282** | WP_005536473.1, EMA21320.1 | 39 | 53 | tafA,B,C,D,E,F,G |
| 52 | **Haloarcula sebkhae JCM 19018** | WP_005536473.1, GGK66820.1 | 39 | 53 | tafA,B,C,D,E,F,G |
| 53 | **Haloarcula salaria JCM 15759** | WP_005536473.1, GGM35199.1 | 39 | 53 | tafA,B,C,D,E,F,G |
| 54 | Haloarcula sinaiiensis ATCC 33800 | WP_004965037.1, EMA11755.1 | 40 | 54 | tafA,B,C,D1?,E,F,G1? + tafD2,G2 |
| 55 | Haloarcula quadrata DSM 11927 | WP_121302043.1, RKS80910.1 - **tafA1** | 39 | 54 | tafA1,B,C1*,D1,E1*,F1,G1,G2 + tafD2,G3 + tafA2C2?E2F2 |
| 56 | Halorubrum terrestre 22517_05_Cabo | WP_159369375.1, MYL17490.1 - **tafA2** | 40 | 55 | tafA1,A2,B1,C,D1,E,F + tafB2,D2,F2 + tafD3,G |
| 57 | **Haloarcula californiae ATCC 33799** | WP_007189710.1, EMA15400.1 | 37 | 53 | tafA,B,C,D1,E,F,G1 + tafD2,G2 + tafD3 |
| 58 | Haloterrigena mahii H13 | OAQ51921.1 | 42 | 56 | tafA,B,D,E,G |
| 59 | Haloarcula sp. JP-Z28 | WP_165897948.1, NHN66102.1 - **tafA1** | 37 | 52 | tafA1,B?,C1,D1,E1,F1,G1,G2 + tafD2,G3 + tafC2,D3,G4 + tatA2E2F2 |
| 60 | Halorubrum sp. ARQ200 | WP_137704928.1, TKX43693.1 - **tafA2** | 35 | 52 | tafA1,C,D1,F + tatA2,D2,E,F + tafB,D3,G |
| 61 | Haloarcula quadrata DSM 11927 | RKS75786.1 - **tafA2** | 36 | 52 | tafA1,B,C1*,D1,E1*,F1,G1,G2 + tafD2,G3 + tafA2C2?E2F2 |
| 62 | **Haloarcula marismortui ATCC 43049** | AAV44314.1 - tafA2 | 36 | 51 | tafA2,C2,E2,G3 (plasmid) + the chromosomal cluster |
| 63 | **Haloarcula californiae ATCC 33799** | WP_080509843.1 | 37 | 54 | tafA,B,C,D1,E,F,G1 + tafD2,G2 + tafD3 |
| 64 | **Haloarcula marismortui ATCC 43049** | WP_011224548.1, AAV47709.1 - **tafA1** | 43 | 55 | tafA1,B,C1,D,E1,F,G1, G2 (chromosome) + the plasmid cluster |
| 65 | Haloarcula sp. JP-Z28 | WP_193569627.1 - **tafA2** | 37 | 54 | tafA1,B?,C1,D1,E1,F1,G1,G2 + tafD2,G3 + tafC2,D3,G4 + tatA2E2F2 |
| 66 | **Haloarcula sp. R1-2** | WP_127013809.1, NHX39577.1 | 38 | 58 | tafA,B,C,D,E,F,G |
| 67 | Natrinema sp. SLN56 | WP_226004493.1 | 38 | 55 | tafA,B,C,D1,E,G1 + tafD2,G2 + tafD3G3 |
| 68 | Salinadaptatus halalkaliphilus XQ-INN 246 | WP_141464248.1, THE65216.1 | 42 | 54 | tafA,C,D,G |
| 69 | **Haloarcula vallismortis ATCC 29715** | EMA00586.1 | 38 | 55 | tafA,B,C,D,E,F,G |
| 70 | Natrialba sp. INN-245 | WP_160047655.1, MWV40343.1 | 36 | 51 | tafA1,A2?,B,C?,D,E?,G |
| 71 | Halorubrum coriense DSM 10284 | ELZ50460.1 - **tafE** | 37 | 53 | tafA,B,D1,E,F + tafD2,G |
| 72 | Natronorubrum thiooxidans HArc-T | WP_143823884.1 | 39 | 51 | tafA,B,C?,D,E,G |
| 73 | Halorubrum sp. WN019 | WP_095638178.1, PAU79996.1 | 42 | 59 | tafA,B,C,D1,E + tafD2,G |
| 74 | **Haloarcula vallismortis ATCC 29715** | WP_004518231.1, EMA00585.1 - **tafE** | 43 | 55 | tafA,B,C,D,E,F,G |
| | * - disrupted by the insertion of the transposase gene. | | | | |

Table 1S

| Protein | Sec Signal Peptide Probability | TAT Signal Peptide Probability | Cleavage Site Position | Cleavage Probability | TwinArg Pattern |
|---|---|---|---|---|---|
| TatA/WP_044951594.1/ HAH_0240 | 0.0363 | 0.9531 | 35-36(VSA-TR) | 0.5960 | NRRSVL |
| TatB/WP_023843080.1/ HAH_0241 | 0.0007 | 0.001 | no | no | no |
| TatC/WP_014039247.1/ HAH_0242 | 0.3869 | 0.531 | 27-28 (TAG-VT) | 0.2381 | TRRGAL |
| TatD/WP_014039248.1/ HAH_0243 | 0.2129 | 0.0096 | no | no | no |
| TatE/WP_044951591.1/ HAH_0239 | 0.0104 | 0.9883 | 31-32 (IEA-DR) | 0.8025 | NRRGFL |
| TatF/WP_023843079.1/ HAH_0238 | 0.9846 | 0.0026 | 21-22 (AMA-TG). | 0.5910 | no |
| TatG/WP_044951588.1/ HAH_0237 | 0.0582 | 0.0049 | no | no | no |

Table 2S

| | TafA | TafB | TafC | TafD | TafE | TafF | TafG | 5NKT |
|---|---|---|---|---|---|---|---|---|
| TafA | Z=36.3, %id=100 | ND | **Z=4.3, %id=19** | **Z=5.7, %id=14** | **Z=8.3, %id=18** | **Z=7.0, %id=9** | ND | **Z=4.5, %id=12** |
| TafB | ND | Z=26.4, %id=100 | ND | **Z=5.6, %id=15** | ND | ND | Z=2.7, %id=15 | ND |
| TafC | **Z=4.3, %id=19** | ND | Z=55.1, %id=100 | **Z=5.1, %id=10** | **Z=4.7, %id=10** | **Z=5.3, %id=18** | **Z=5.4, %id=13** | Z=3.6, %id=9 |
| TafD | **Z=5.7, %id=14** | **Z=5.6, %id=15** | **Z=5.1, %id=10** | Z=52.6, %id=100 | Z=3.6, %id=5 | **Z=5.2, %id=11** | Z=3.6, %id=7 | Z=3.8, %id=8 |
| TafE | **Z=8.3, %id=18** | ND | **Z=4.7, %id=10** | Z=3.6, %id=5 | Z=33.6, %id=100 | **Z=10.0, %id=16** | ND | **Z=6.8, %id=15** |
| TafF | **Z=7.0, %id=9** | ND | **Z=5.3, %id=18** | **Z=5.2, %id=11** | **Z=10.0, %id=16** | Z=33.0, %id=100 | Z=2.5, %id=12 | **Z=5.4, %id=13** |
| TafG | ND | Z=2.7, %id=15 | **Z=5.4, %id=13** | Z=3.6, %id=7 | ND | Z=2.5, %id=12 | Z=38.0, %id=100 | ND |
| 5NKT | **Z=4.5, %id=12** | ND | Z=3.6, %id=9 | Z=3.8, %id=8 | **Z=6.8, %id=15** | **Z=5.4, %id=13** | ND | Z=31.6, %id=100 |

Table 3S

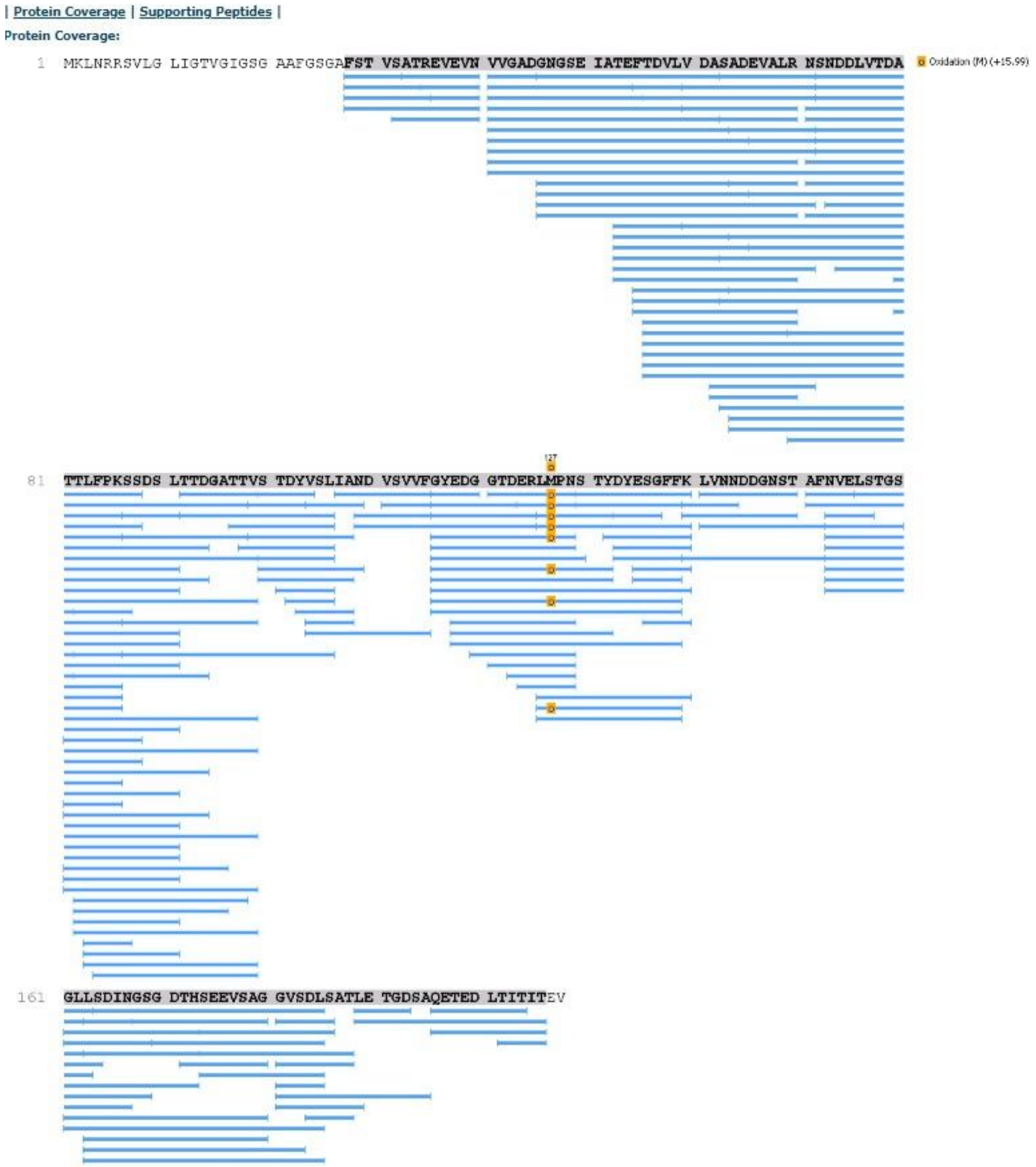
| Protein Coverage | Supporting Peptides |
Protein Coverage:
1 MKLNRRSVLG LIGTVGIGSG AAFGSGAFST VSATREVEVN VVGADGNGSE IATEFTDVLV DASADEVALR NSNDDLVTDA
Oxidation (M) (+15.99)
127
81 TTLFPKSSDS LTTDGATTVS TDYVSLIAND VSVVFGYEDG GTDERLMPNS TYDYESGFFK LVNNDDGNST AFNVELSTGS
161 GLLSDINGSG DTHSEEVSAG GVSDLSATLE TGDSAQETED LTITITEV

F1S

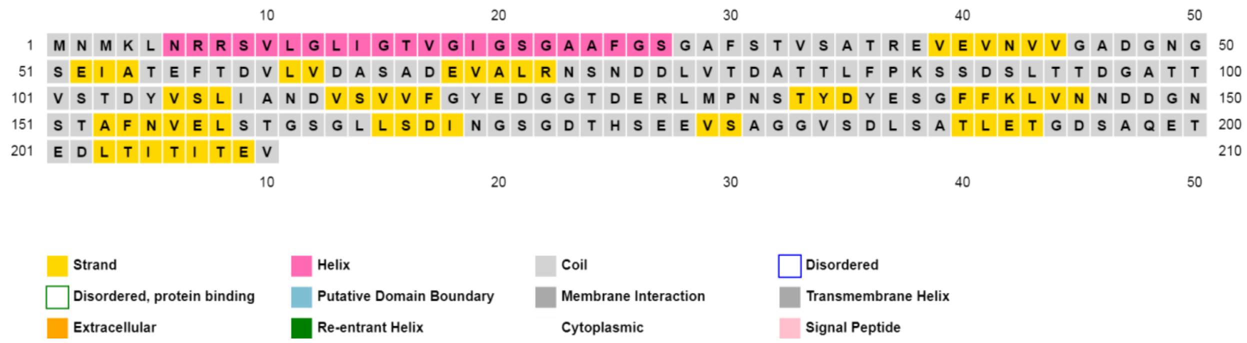
10 20 30 40 50
1 M N M K L N R R S V L G L I G T V G I G S G A A F G S G A F S T V S A T R E V E V N V V G A D G N G 50
51 S E I A T E F T D V L V D A S A D E V A L R N S N D D L V T D A T T L F P K S S D S L T T D G A T T 100
101 V S T D Y V S L I A N D V S V V F G Y E D G G T D E R L M P N S T Y D Y E S G F F K L V N N D D G N 150
151 S T A F N V E L S T G S G L L S D I N G S G D T H S E E V S A G G V S D L S A T L E T G D S A Q E T 200
201 E D L T I T I T E V 210
10 20 30 40 50
Strand
Helix
Coil
Disordered
Disordered, protein binding
Putative Domain Boundary
Membrane Interaction
Transmembrane Helix
Extracellular
Re-entrant Helix
Cytoplasmic
Signal Peptide

**TafA**

F2S

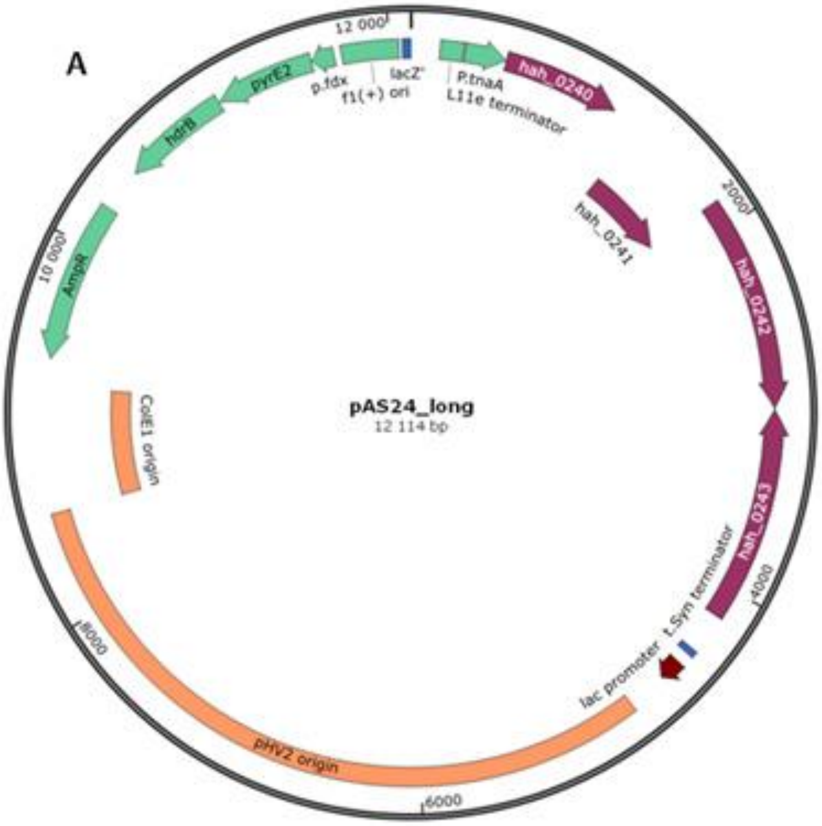
A
12 000
p.fdx
lacZ'
f1(+) ori
P.tnaA
L11e terminator
hah_0240
pyrE2
hdrB
hah_0241
2000
hah_0242
10 000
AmpR
pAS24_long
12 114 bp
ColE1 origin
hah_0243
t.Syn terminator
4000
8000
lac promoter
pHV2 origin
6000

F3S

### Band 1:

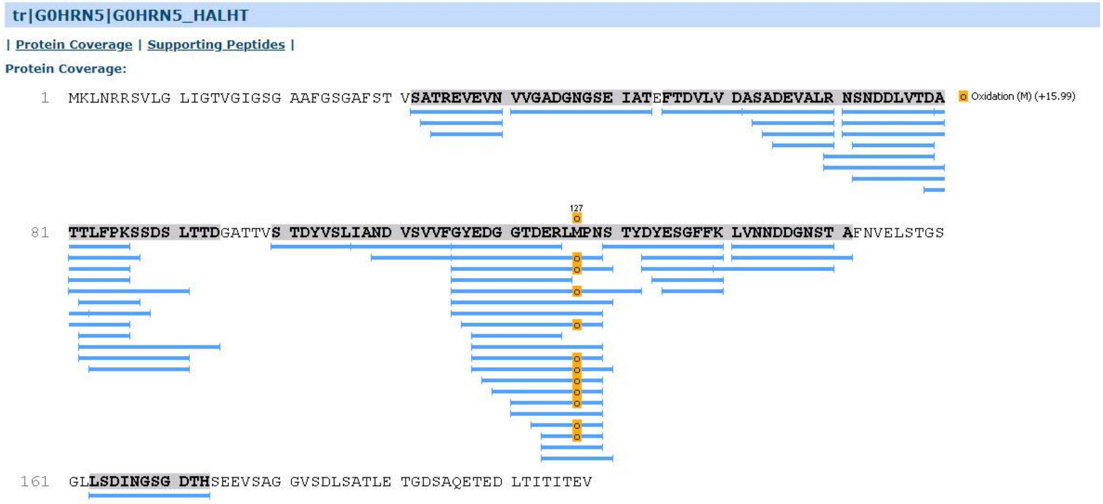

### TafA

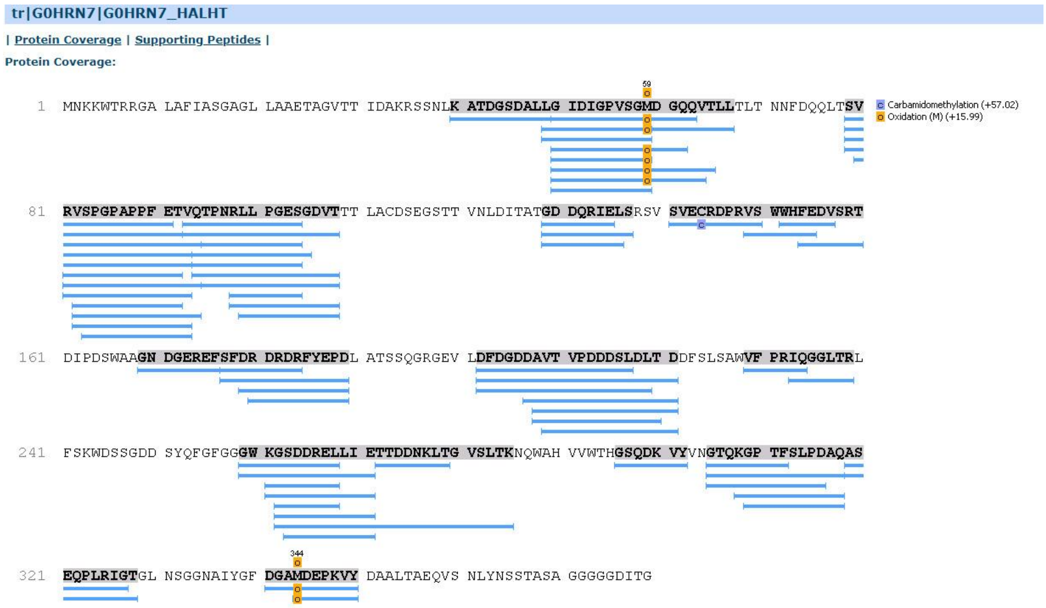

### TafC

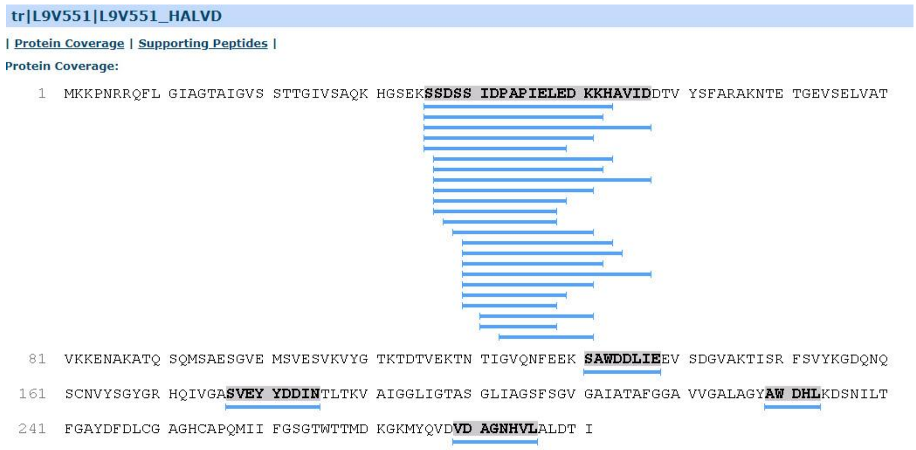

### HVO_A0133

### Band 2:

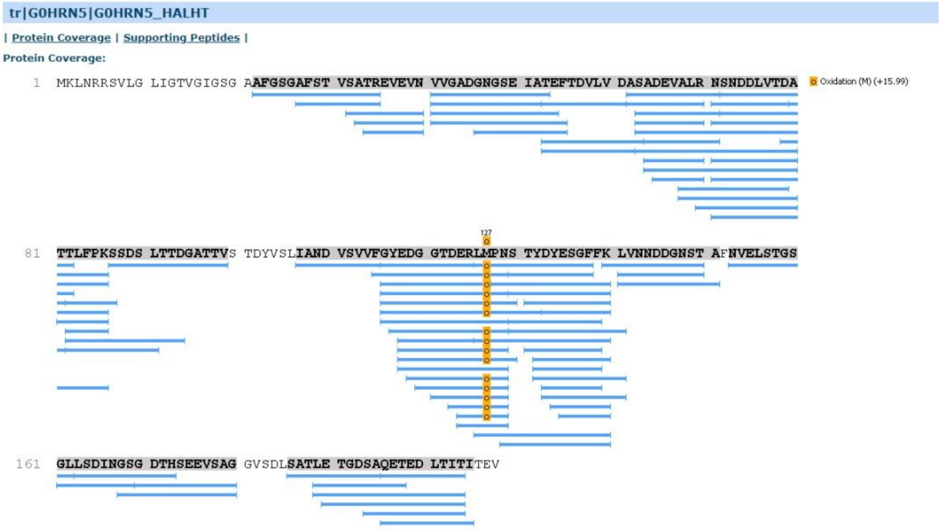

#### TafA

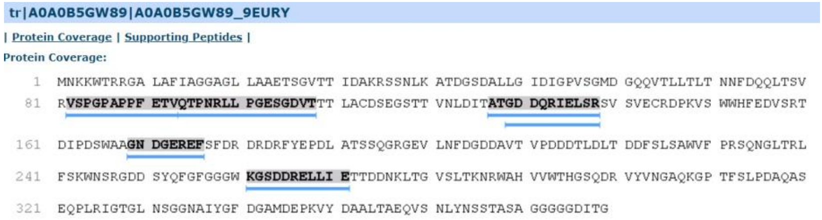

#### TafC

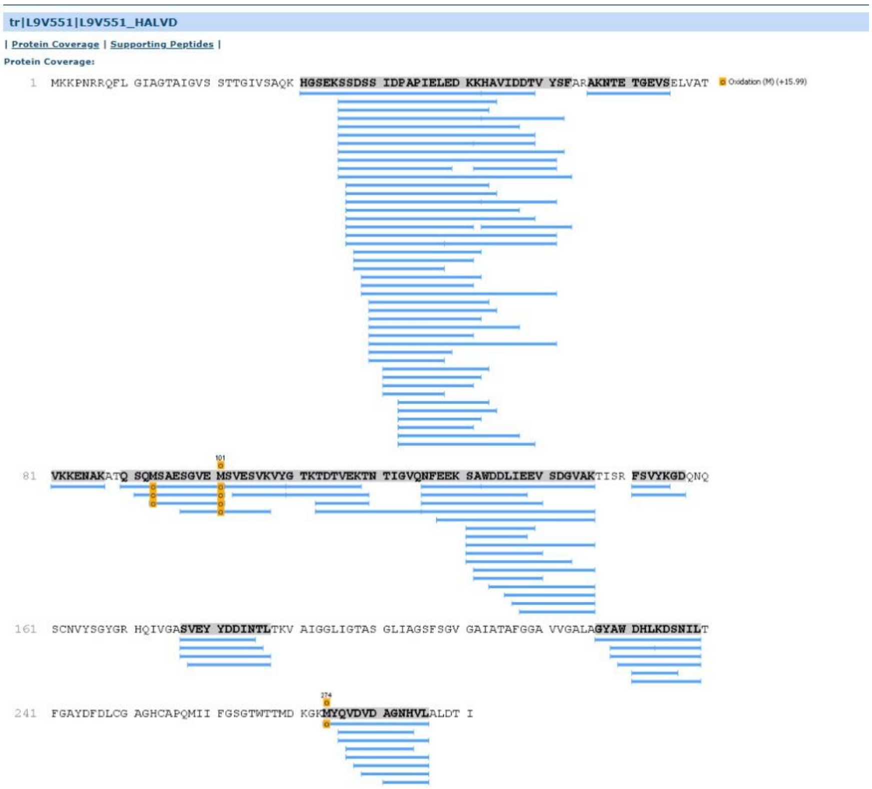

#### HVO_A0133

### Band 3:

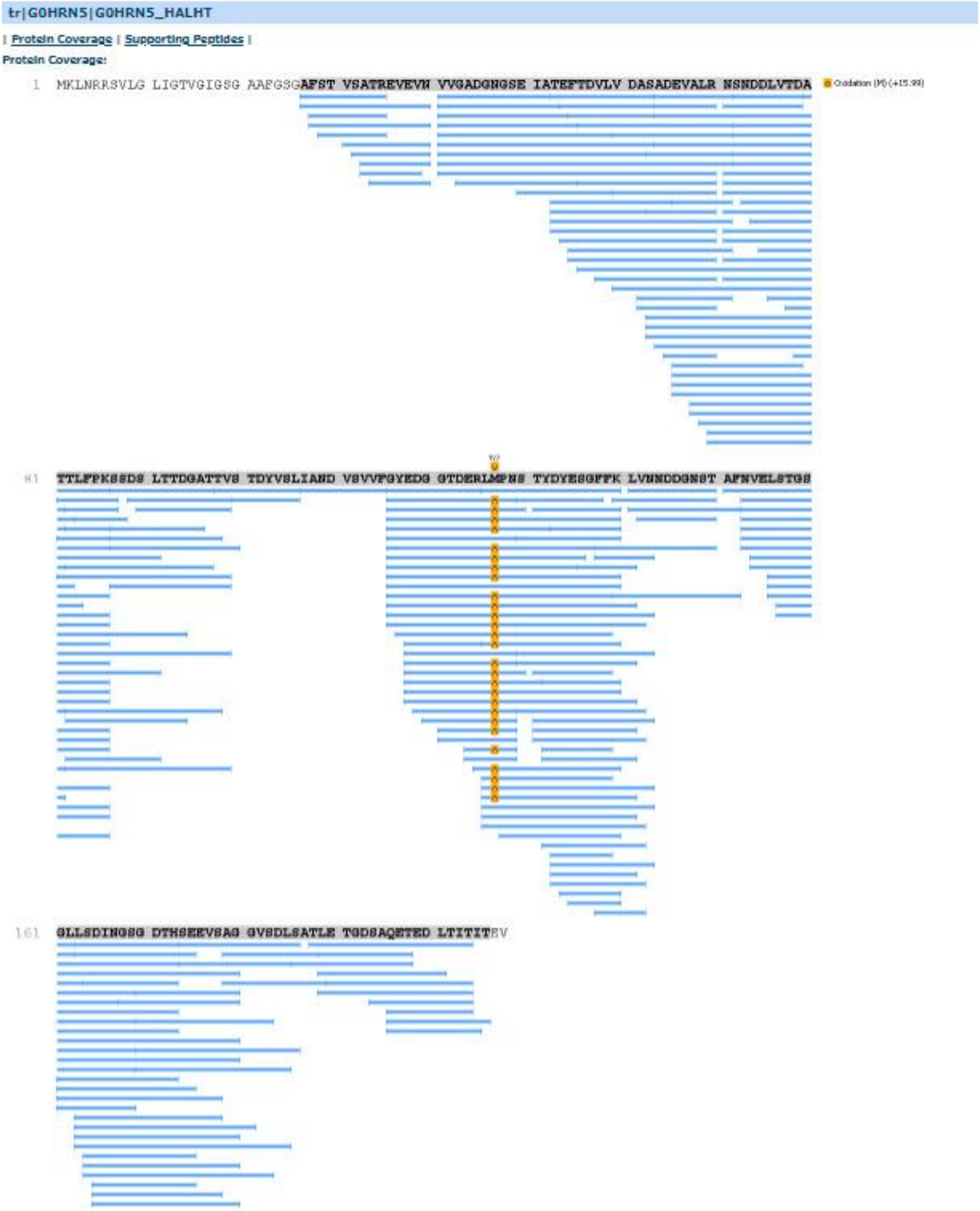

### TafA

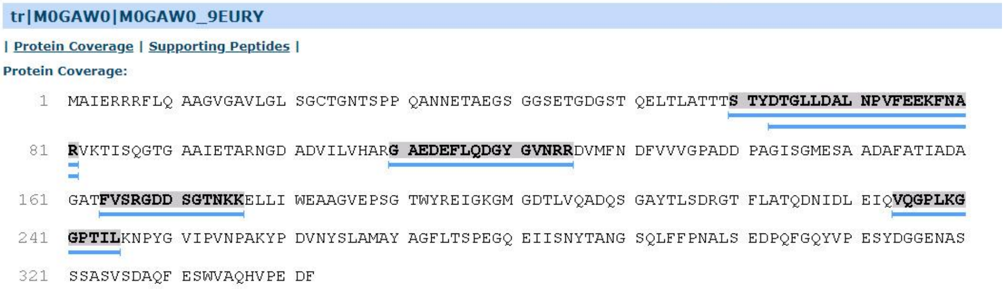

### TafC

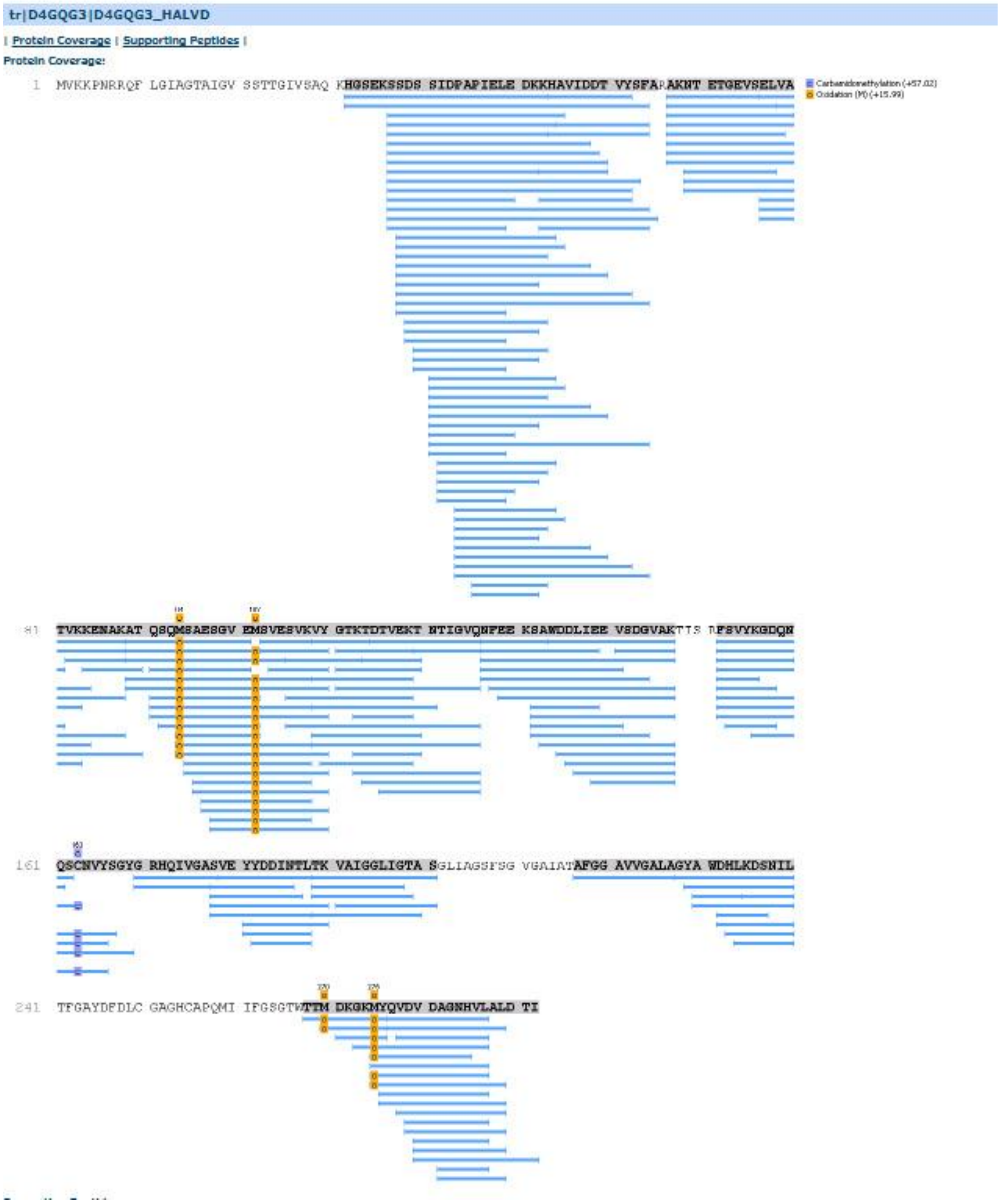

### HVO_A0133

### Band 4:

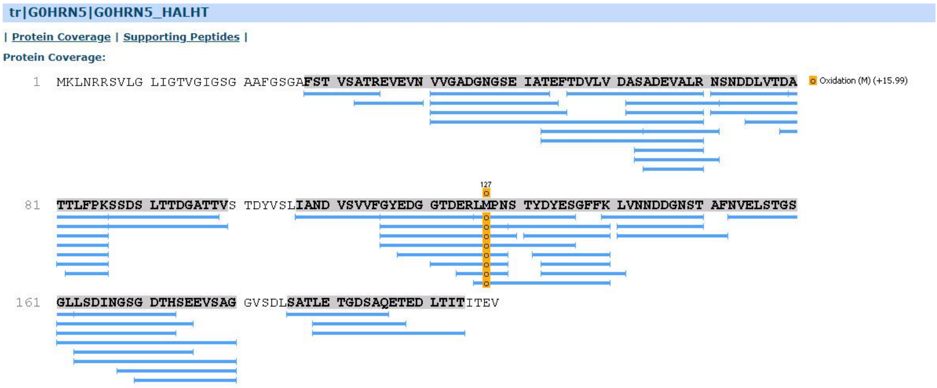

### TafA

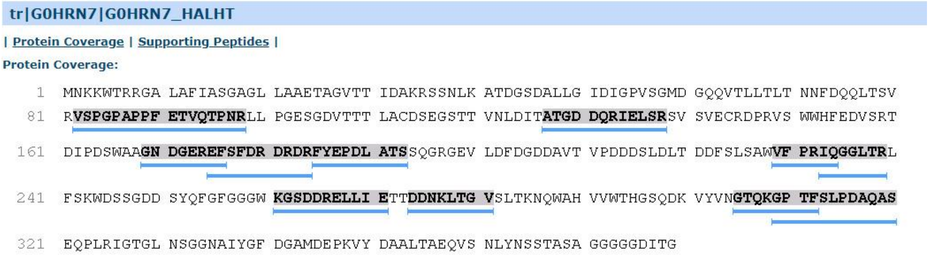
tr|G0HRN7|G0HRN7_HALHT
| Protein Coverage | Supporting Peptides |
Protein Coverage:
1 MNKKWTRRGA LAFIASGAGL LAAETAGVTT IDAKRSSNLK ATDGSDALLG IDIGPVSGMD GQQVTLLTLT NNFDQQLTSV
81 RVSPGPAPPF ETVQTPNRLL PGESGDVTTT LACDSEGSTT VNLDITATGD DQRIELSRSV SVECRDPRVS WWHFEDVSRT
161 DIPDSWAAGN DGEREFSFDR DRDRFYEPDL ATSSQGRGEV LDFDGDDAVT VPDDDSLDLT DDFSLSAWVF PRIQGGLTRL
241 FSKWDSSGDD SYQFGFGGGW KGSDDRELLI ETTDDNKLTG VSLTKNQWAH VVWTHGSQDK VYVNGTQKGP TFSLPDAQAS
321 EQPLRIGTGL NSGGNAIYGF DGAMDEPKVY DAALTAEQVS NLYNSSTASA GGGGGDITG

### TafC

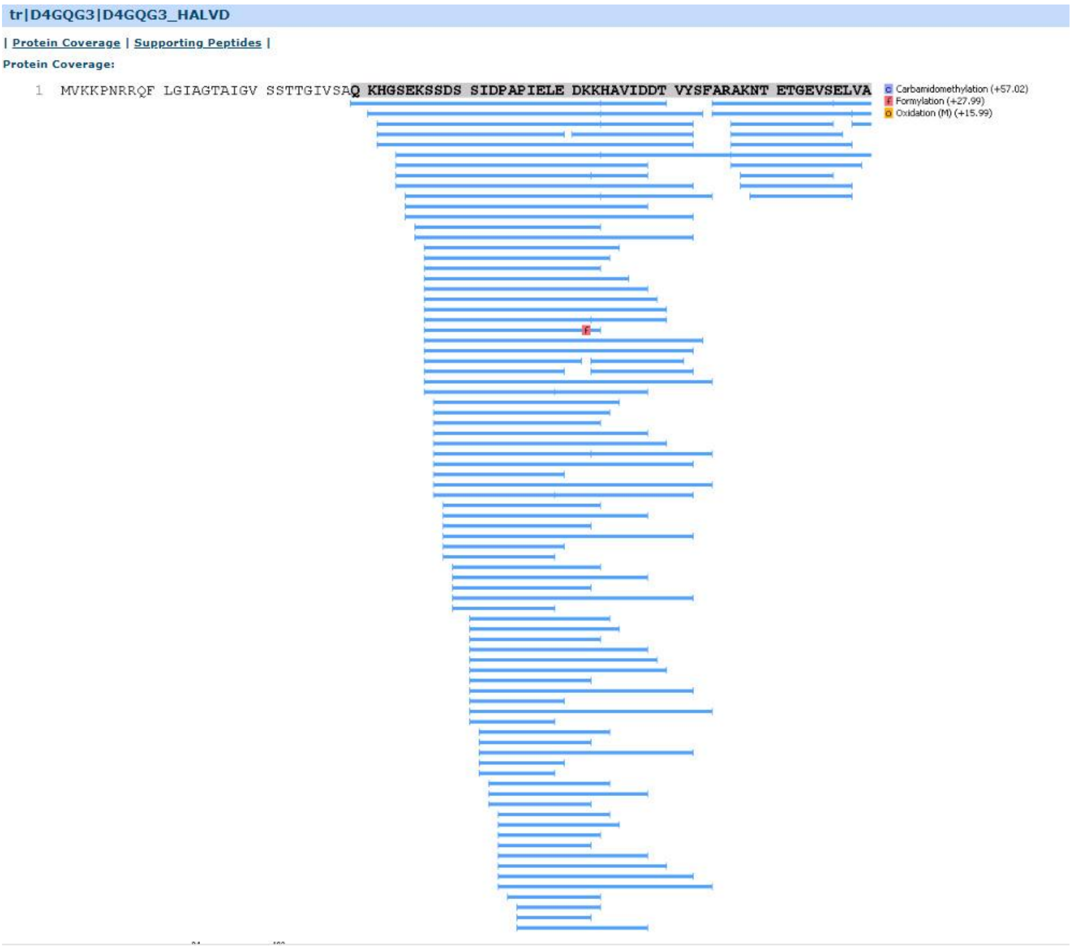
tr|D4GQG3|D4GQG3_HALVD
| Protein Coverage | Supporting Peptides |
Protein Coverage:
1 MVKKPNRRQF LGIAGTAIGV SSTTGIVSAQ KHGSEKSSDS SIDPAPIELE DKKHAVIDDT VYSFARAKNT ETGEVSELVA
Carbamidomethylation (+57.02)
Formylation (+27.99)
Oxidation (M) (+15.99)

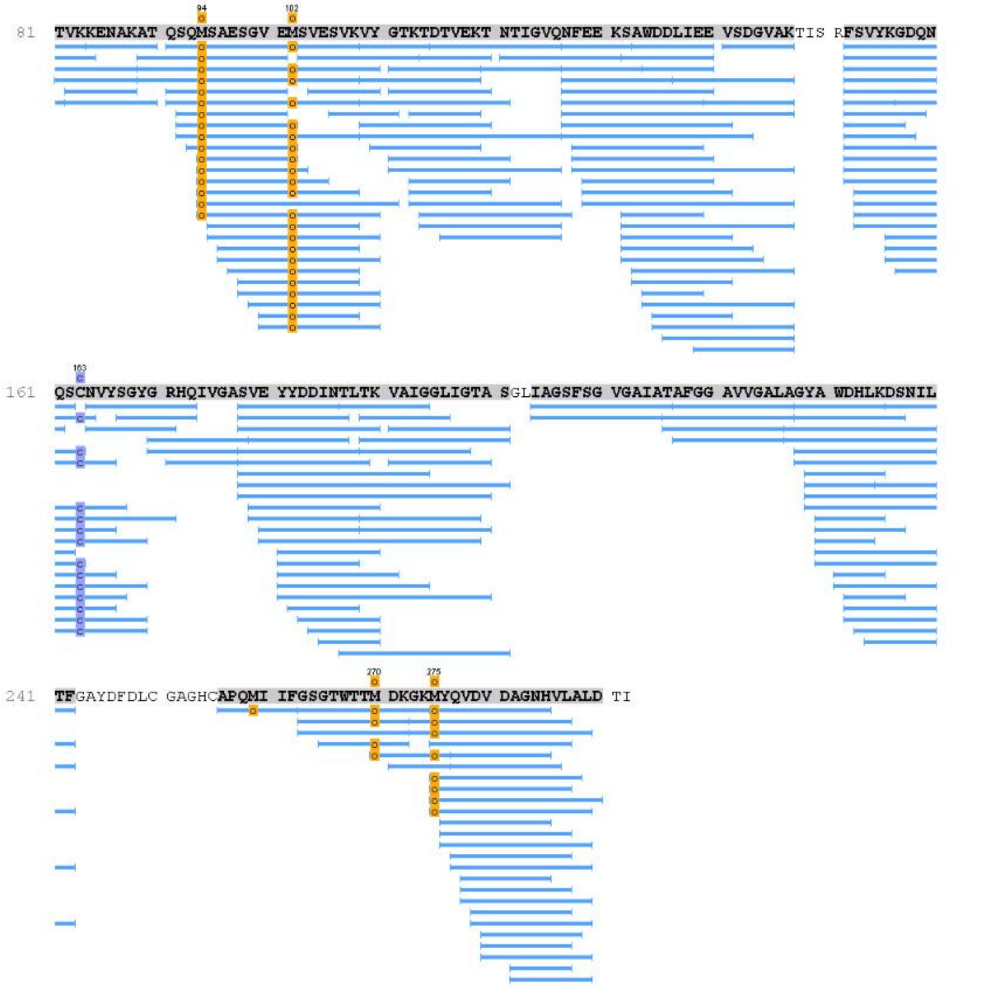

**HVO_A0133**

F4S

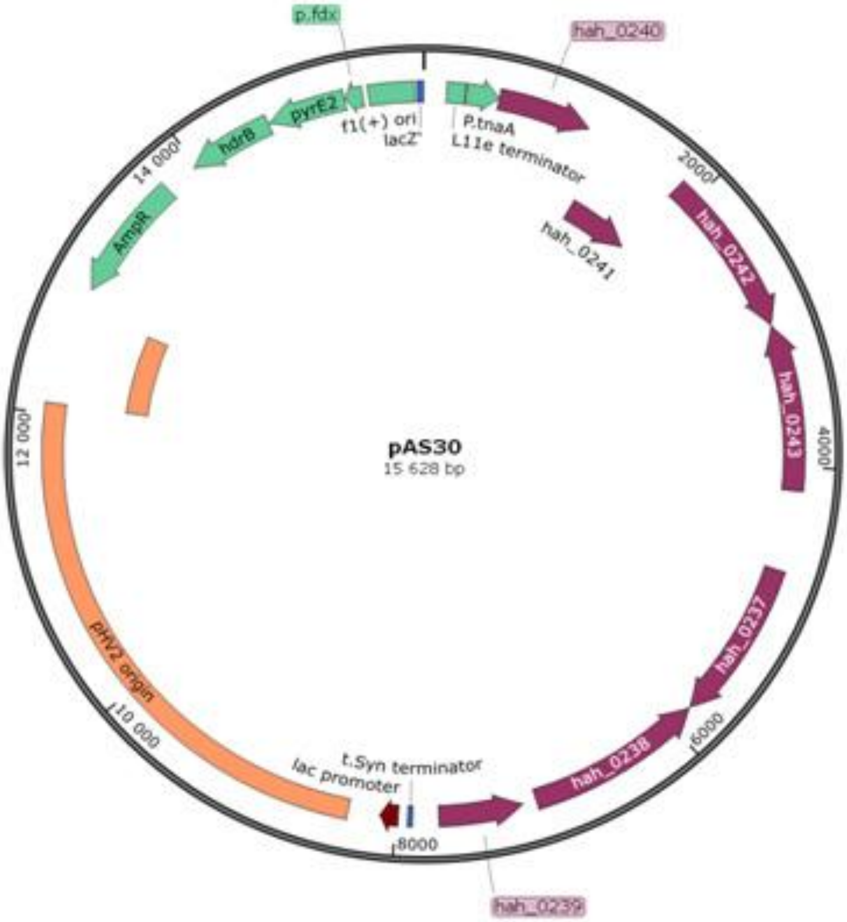
p.fdx
hah_0240
f1(+) ori
lacZ'
P.tnaA
L11e terminator
pyrE2
hdrB
14 000
2000
AmpR
hah_0241
hah_0242
hah_0243
pAS30
15 628 bp
12 000
4000
hah_0237
pHV2 origin
10 000
6000
hah_0238
t.Syn terminator
lac promoter
8000
hah_0239

F5S

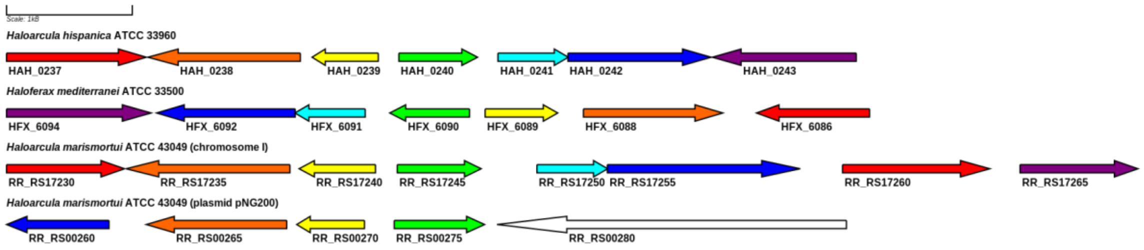
Scale: 1kB
Haloarcula hispanica ATCC 33960
HAH_0237
HAH_0238
HAH_0239
HAH_0240
HAH_0241
HAH_0242
HAH_0243
Haloferax mediterranei ATCC 33500
HFX_6094
HFX_6092
HFX_6091
HFX_6090
HFX_6089
HFX_6088
HFX_6086
Haloarcula marismortui ATCC 43049 (chromosome I)
RR_RS17230
RR_RS17235
RR_RS17240
RR_RS17245
RR_RS17250
RR_RS17255
RR_RS17260
RR_RS17265
Haloarcula marismortui ATCC 43049 (plasmid pNG200)
RR_RS00260
RR_RS00265
RR_RS00270
RR_RS00275
RR_RS00280

F6S

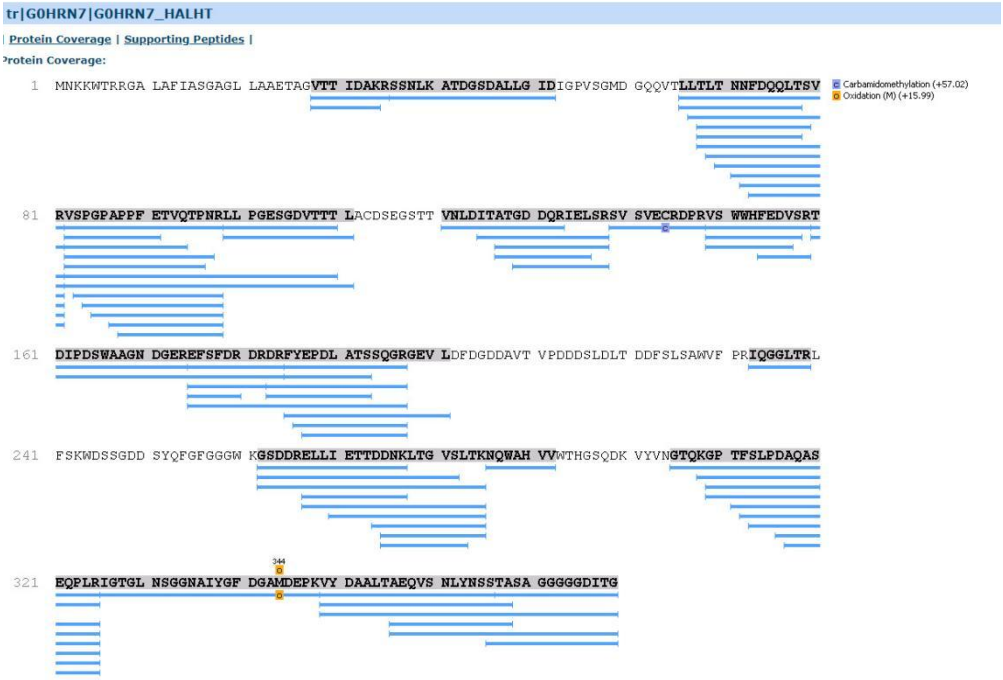
tr|G0HRN7|G0HRN7_HALHT
| Protein Coverage | Supporting Peptides |
rotein Coverage:
1 MNKKWTRRGA LAFIASGAGL LAAETAGVTT IDAKRSSNLK ATDGSDALLG IDIGPVSGMD GQQVTLLTLT NNFDQQLTSV
Carbamidomethylation (+57.02)
Oxidation (M) (+15.99)
81 RVSPGPAPPF ETVQTPNRLL PGESGDVTTT LACDSEGSTT VNLDITATGD DQRIELSRSV SVECRDPRVS WWHFEDVSRT
161 DIPDSWAAGN DGEREFSFDR DRDRFYEPDL ATSSQGRGEV LDFDGDDAVT VPDDDSLDLT DDFSLSAWVF PRIQGGLTRL
241 FSKWDSSGDD SYQFGFGGGW KGSDDRELLI ETTDDNKLTG VSLTKNQWAH VVWTHGSQDK YVYNGTQKGP TFSLPDAQAS
344
321 EQPLRIGTGL NSGGNAIYGF DGAMDEPKVY DAALTAEQVS NLYNSSTASA GGGGGDITG

**A**

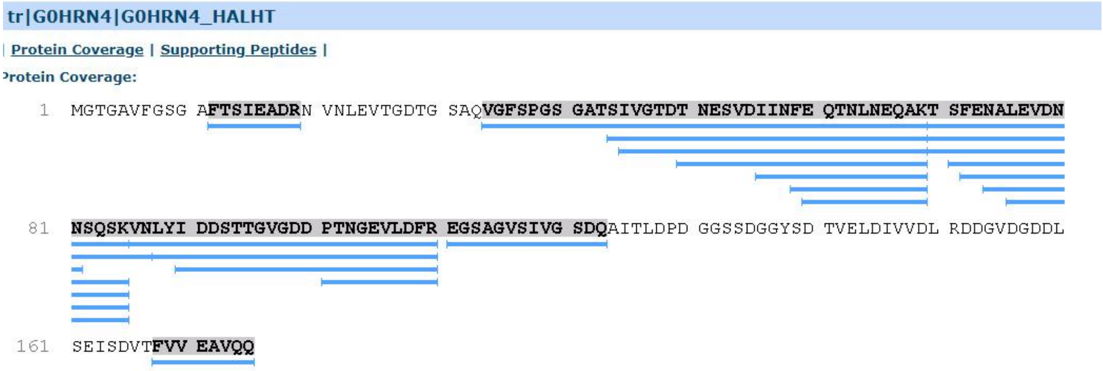
tr|G0HRN4|G0HRN4_HALHT
| Protein Coverage | Supporting Peptides |
rotein Coverage:
1 MGTGAVFGSG AFTSIEADRN VNLEVTGDTG SAQVGFSPGS GATSIVGTDT NESVDIINFE QTNLNEQAKT SFENALEVDN
81 NSQSKVNLYI DDSTTGVGDD PTNGEVLDFR EGSAGVSIVG SDQAITLDPD GGSSDGGYSD TVELDIVVDL RDDGVDGDDL
161 SEISDVTFVV EAVQQ

**B**

F7S
